# Structural basis for the broad recognition specificity of an Arabidopsis immune receptor

**DOI:** 10.64898/2026.02.04.703898

**Authors:** He Zhao, Camille-Madeleine Szymansky, Xue Lyu, Jianhua Huang, Paul Derbyshire, Frank L H Menke, Michael W Webster, Sophien Kamoun, Muniyandi Selvaraj, Jonathan D G Jones

## Abstract

Plant nucleotide-binding leucine-rich repeat (NLR) immune receptors typically confer resistance through recognition of specific pathogen effectors. The Arabidopsis NLR WRR4A defies this paradigm by recognizing multiple sequence-divergent effectors from *Albugo candida*, conferring resistance to multiple pathogen races. Despite minimal sequence similarity, these effectors share a conserved N-terminal ferredoxin-like fold. Through cryo-EM structure determination of two WRR4A resistosomes bound to sequence-distinct effectors, combined with AlphaFold modelling, we reveal a shape-based recognition mechanism: WRR4A engages structurally conserved backbone features of the effectors in a mostly side chain-independent manner, enabling recognition of diverse effectors with similar three-dimensional architectures. These insights guided successful engineering of WRR4A to acquire novel recognition specificity. In addition, analysis of the monomeric WRR4A resting state reveals a distinct domain architecture characteristic of C-JID–containing TIR-NLRs and informs their activation mechanism. This work provides insights into NLR-mediated broad-spectrum recognition and the potential for structure-informed engineering of improved crop resistance.

## INTRODUCTION

NLRs are intracellular immune receptors defined by their characteristic nucleotide-binding domain (NBD) and leucine-rich repeats (LRRs). In three kingdoms of life, they detect non-self or modified-self molecules and trigger defense responses, constituting a crucial part of the innate immune system(Jones et al., 2016; Meunier and Broz, 2017; Kibby et al., 2023). Mammalian NLRs primarily detect conserved molecular patterns associated with pathogen invasion and typically exist in relatively small repertoires. Plant NLRs, by contrast, have evolved to recognize strain-specific effectors (virulence factors delivered from pathogens) that exhibit substantial variation across pathogen species and strains(Maekawa et al., 2011). To counter this diversity, plants have evolved extensive repertoires of highly polymorphic NLRs(Maekawa et al., 2011; Barragan and Weigel, 2021). The coevolutionary dynamics between NLRs and pathogen effectors shape plant-pathogen interactions and elucidating the molecular basis of receptor recognition and pathogen evasion is essential for understanding this coevolution(Zhang et al., 2022; Contreras et al., 2023; Jones et al., 2024).

Specific effector recognition by plant NLRs underpins the gene-for-gene model, in which a plant resistance gene detects its cognate pathogen effector to activate defense(Flor, 1971). Although NLRs have been widely exploited in crop breeding to reduce yield losses, this high specificity frequently renders NLR-mediated resistance short-lived in the field. Under selection pressure imposed by a single NLR, pathogen populations can rapidly evade recognition through loss, mutation, or modification of the recognized effector, outpacing the evolutionary capacity of host NLRs(Dangl et al., 2013). In theory, NLRs with broader recognition spectra are expected to confer more durable resistance, yet such mechanisms have rarely been documented. An alternative strategy is the deployment of multiple NLRs within a single cultivar, known as NLR stacking, although identifying compatible NLR combinations remains challenging(Dangl et al., 2013; Luo et al., 2021; Zhao et al., 2025). In this context, structure-guided engineering of NLRs offers a promising avenue for generating novel and potentially broader recognition specificities(Liu et al., 2021; Kourelis et al., 2023; Wang et al., 2025), provided that sufficient structural and mechanistic information is available. Recent advances in structural biology have yielded cryo-EM structures of several plant NLRs with their cognate effectors (ZAR1(Wang et al., 2019a; Wang et al., 2019b), Sr35(Forderer et al., 2022), ROQ1(Martin et al., 2020), RPP1(Ma et al., 2020), and MLA13(Lawson et al., 2025)), revealing diverse recognition mechanisms. The coiled-coil (CC)-NLR ZAR1 indirectly recognizes AvrAC through PBL2 kinase modification, while other CC-NLRs like Sr35 and MLA13 directly bind effectors primarily through their LRR domains. ROQ1 and RPP1, which belong to the Toll/interleukin-1 receptor/resistance protein (TIR)-NLR class, feature an extended C-terminal jelly roll/Ig-like domain (C-JID) following their LRR domains and engage effectors through both LRR and C-JID domains(Ma et al., 2020; Martin et al., 2020). Using these structural insights, researchers have exchanged recognition capabilities between natural alleles of Sr35 and MLA13(Lawson et al., 2025). However, engineering entirely novel recognition specificities using structural information alone, without depending on evolutionary patterns or natural variants, remains challenging.

The TIR-NLR WRR4A (WHITE RUST RESISTANCE 4) from the Arabidopsis Col-0 ecotype confers resistance against several races of *Albugo candida*, an oomycete pathogen that causes white blister rust, a significant disease of Brassica crops(Borhan et al., 2010). We previously defined the genome of isolate Ac2V and characterized more than one hundred effectors containing a conserved “Cx₂Cx₅G” (CCG) motif(Furzer et al., 2022). Unusually, WRR4A can recognize at least eight CCG effectors through their N-terminal ∼100 amino acids, which likely explains the broad-spectrum resistance that WRR4A provides against most *A. candida* races(Redkar et al., 2023). Remarkably, the recognized regions from different CCG effectors share less than 20% sequence identity: a level of divergence that would typically abolish recognition in other characterized NLR recognition systems. This broad recognition capacity makes the WRR4A-CCG recognition system an exceptional model for understanding how a single NLR can recognize multiple sequence-distinct effectors.

We investigated the molecular basis of the broad recognition capacity of WRR4A by determining cryo-EM structures of WRR4A bound to two sequence-distinct CCG effectors. Comparative analysis indicates that these ferredoxin-like effectors adopt similar bound conformations, contacting the WHD, LRR, and C-JID domains of WRR4A, with LRR and C-JID as the primary interfaces. Integration of experimental structures with AlphaFold predictions further reveals that, despite sequence divergence, WRR4A recognizes the conserved structural signature of the N-terminal ferredoxin fold across CCGs, primarily through backbone-mediated interactions at defined structural positions rather than side-chain identity. Guided by these insights, we successfully engineered WRR4A to gain new recognition. Furthermore, structural analysis of the WRR4A resting (resting) state revealed a distinct monomeric architecture in which the NB domain lies against the LRR concave face, contacting both LRR and C-JID in a manner that resembles effector engagement. We propose that effector binding repositions the NB domain, exposing oligomerization interfaces and enabling resistosome assembly. This resting state architecture may be shared among C-JID-containing TIR-NLRs and enabled generation of a sensitized WRR4A variant activated by lower effector concentrations. Overall, our work demonstrates how shape-based recognition enables an NLR to tolerate effector sequence diversification and highlights the potential of structure-guided NLR engineering.

## RESULTS

### WRR4A recognizes conserved N-terminal ferredoxin-like folds of sequence-divergent CCG effectors

WRR4A recognizes eight *Albugo candida* CCG effectors through interactions with their N-terminal ∼100 residues(Redkar et al., 2023). To elucidate the molecular basis of this broad recognition spectrum, we compared the amino acid sequences of recognized and unrecognized CCGs. Since the N-terminal domain alone mediates recognition, we focused our alignment on this region. Among the recognized CCGs, CCG28N, CCG30N, and CCG71N share the highest sequence identity (∼40%), suggesting they may be recognized through similar sequence features. Surprisingly, the remaining recognized CCG members exhibit low sequence identity (13-20%) among their N-terminal domains, comparable to that observed across all CCGs in the table (Fig. 1A, Fig. S1A). This unexpected pattern indicates that WRR4A recognition extends beyond simple sequence conservation. Adding to the complexity, alanine substitutions of the highly conserved cysteine and glycine residues within the CCG do not abolish WRR4A-mediated recognition(Redkar et al., 2023).

**Fig. 1.**
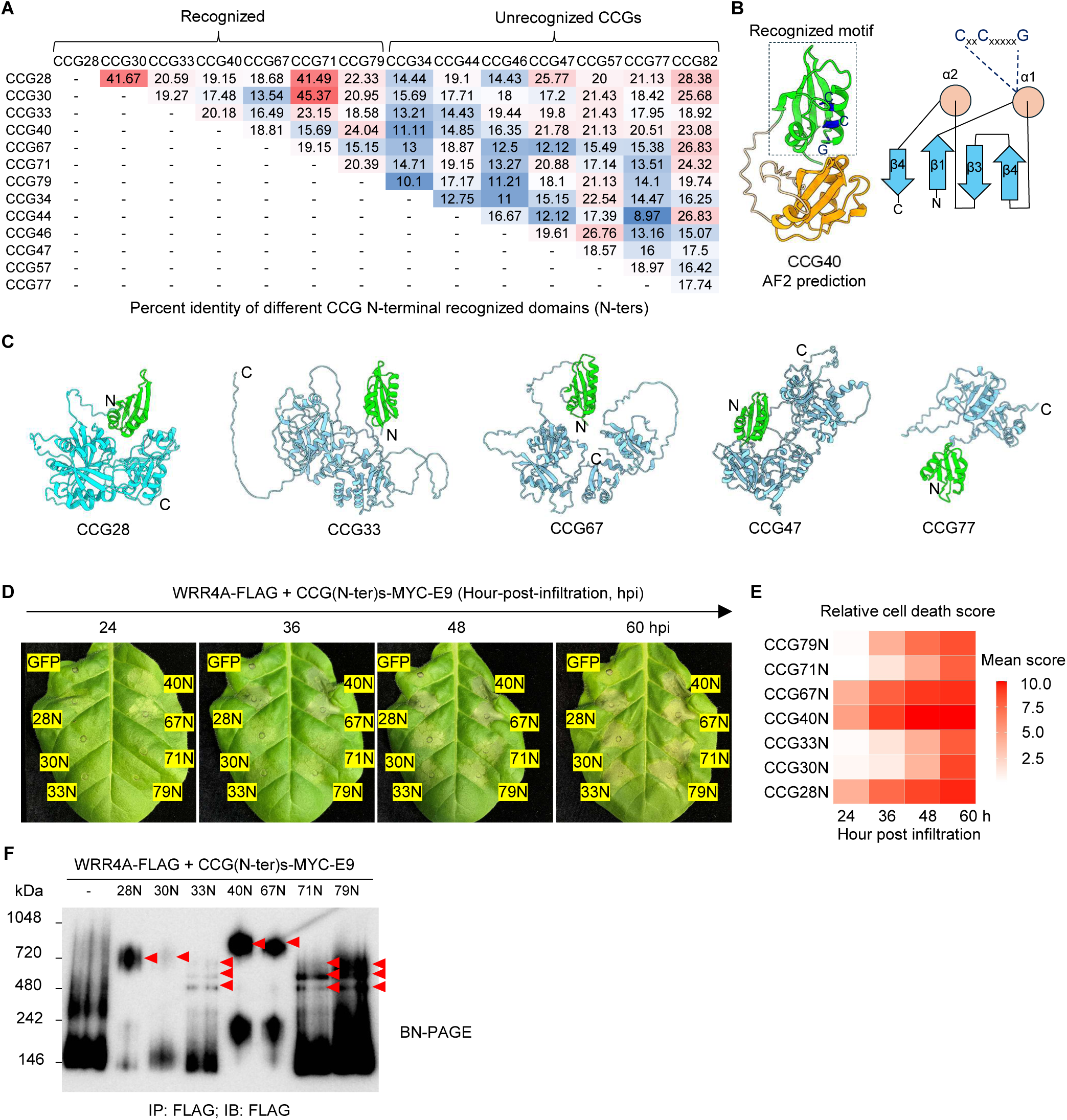
WRR4A recognizes sequence-divergent, structurally conserved N-terminal ferredoxin-like folds of CCG effectors. **(A)** Pairwise sequence identity of the N-terminal ∼100 amino acids of both representative WRR4A-recognized and unrecognized CCG effectors. Signal peptides were removed. Sequences were aligned using MUSCLE. **(B)** AlphaFold2-predicted structure of CCG40. The N-terminal WRR4A-recognized domain (left, labelled in green) adopts a ferredoxin-like fold characterized by a conserved βαββαβ secondary-structure topology (right). **(C)** AlphaFold2 models of representative CCG effectors after removing signalling peptides. **(D and E)** Time-course analysis of WRR4A-mediated cell death triggered by CCG N-terminal domains. MYC-E9-tagged CCG N-terminal regions were co-expressed with WRR4A-FLAG in *Nicotiana tabacum*. A reduced Agrobacterium concentration (each of OD_600_ = 0.1) was used to slow cell death progression and enable clearer discrimination between different CCGs. Data represent means from five infiltrated leaves. **(F)** Oligomerization of WRR4A induced by different CCG N-terminal domains analyzed by blue-native PAGE (BN-PAGE). Red arrows indicate potential WRR4A oligomers.

To investigate how WRR4A recognizes diverse CCG sequences, we predicted the 3D structures of CCGs to assess their structural similarity. Comparison of AlphaFold2(Jumper et al., 2021) models across different CCGs revealed that all are multi-domain proteins. Remarkably, despite low sequence identity, all CCGs share a conserved N-terminal fold (Fig. 1B, C), which is connected via a long, flexible loop to a comparatively divergent C-terminal region of variable length. Notably, this conserved N-terminal fold corresponds precisely to the recognition domain (N-terminal 100 residues) and is fully solvent-exposed, suggesting that WRR4A recognizes this structural feature. Further analysis revealed that this N-terminal fold adopts a βαββαβ secondary structure (Fig. 1B right) - a signature of the ferredoxin-like fold found across multiple protein families with diverse functions. The conserved CCG motif localizes to the α1 helix (Fig. 1B). The ferredoxin-like fold belongs to a class of superfolds that has diversified extensively during evolution(Nishina et al., 2022). Structural searches using Foldseek(van Kempen et al., 2024) revealed that other plant pathogens, including *Magnaporthe oryzae*, *Colletotrichum gloeosporioides*, and *Venturia inaequalis*, also encode secreted proteins with this fold, indicating that ferredoxin-like fold effectors may be widespread among plant pathogens (Fig. S1B).

Previously, WRR4A-recognized CCGs were identified through end-point hypersensitive response (HR) assays, in which all recognizable CCGs triggered comparable levels of cell death when co-expressed with WRR4A in *Nicotiana tabacum(Redkar et al., 2023)*. However, given their sequence variability, we hypothesized that recognition strength could differ among CCGs. We conducted a semi-quantitative time-course HR experiment, monitoring cell death every 12 hours following co-infiltration of WRR4A with the N-terminal recognition domain of different CCGs. While all CCG N-termini triggered similar cell death phenotypes at the final timepoint (60 hpi), they induced distinct levels of cell death at early stages, with CCG28N, CCG40N, and CCG67N eliciting the strongest HR (Fig. 1D, E), suggesting that these three effectors are more efficiently recognized by WRR4A. Consistent with the time-course cell death results, Blue-native PAGE further revealed that the N-termini of CCG28, CCG40, and CCG67—despite sharing only 18-20% sequence identity—induced the most robust WRR4A oligomerization (Fig. 1F). We selected CCG28N, CCG40N and CCG67N as candidates for structural studies.

### Molecular architecture of WRR4A-CCG40N resistosome

To elucidate the structural basis of WRR4A-mediated recognition, we co-expressed FLAG-tagged WRR4A with Strep-MBP-tagged N-terminal recognized domains of CCG28, CCG40, and CCG67 separately in *N. benthamiana eds1* mutants(Schultink et al., 2017), in which TIR-mediated cell death is abolished. The WRR4A-CCG complexes were purified by sequential Strep-Tactin and FLAG affinity purification for cryo-EM structure determination. Although comparable amounts of protein were obtained for all three samples (Fig. S1C), negative-stain EM revealed that purified WRR4A-CCG40N and WRR4A-CCG67N complexes were more homogeneous than WRR4A-CCG28N, displaying uniform four-leaf clover structures resembling the TIR-NLRs RPP1- and ROQ1- resistosomes(Ma et al., 2020; Martin et al., 2020) (Fig. 2A). In contrast, the WRR4A-CCG28N sample contained additional small particles alongside the large tetramers, indicating potential degradation or complex instability (Fig. 2A). Activated TIR-NLR resistosomes function as NAD^+^ hydrolases(Ma et al., 2020). All three WRR4A-CCG complexes exhibited NADase activity when incubated with the NAD^+^ fluorescent analog ε-NAD, confirming successful purification of functional WRR4A resistosomes (Fig. 2B). Despite its heterogeneity, the WRR4A-CCG28N sample displayed NADase activity comparable to the other complexes (Fig. 2B). While some TIR-only proteins and isolated TIR domains from full-length TIR-NLRs exhibit nuclease activity(Yu et al., 2022), neither the WRR4A resistosome nor resting state WRR4A showed detectable RNA-hydrolyzing activity compared with the TIR-domain protein RBA1 (Fig. S1D). Conceivably, full-length TIR-NLRs may lack nuclease activity due to an inability to assemble the filamentous architectures required for DNA/RNA hydrolysis.

**Fig. 2.**
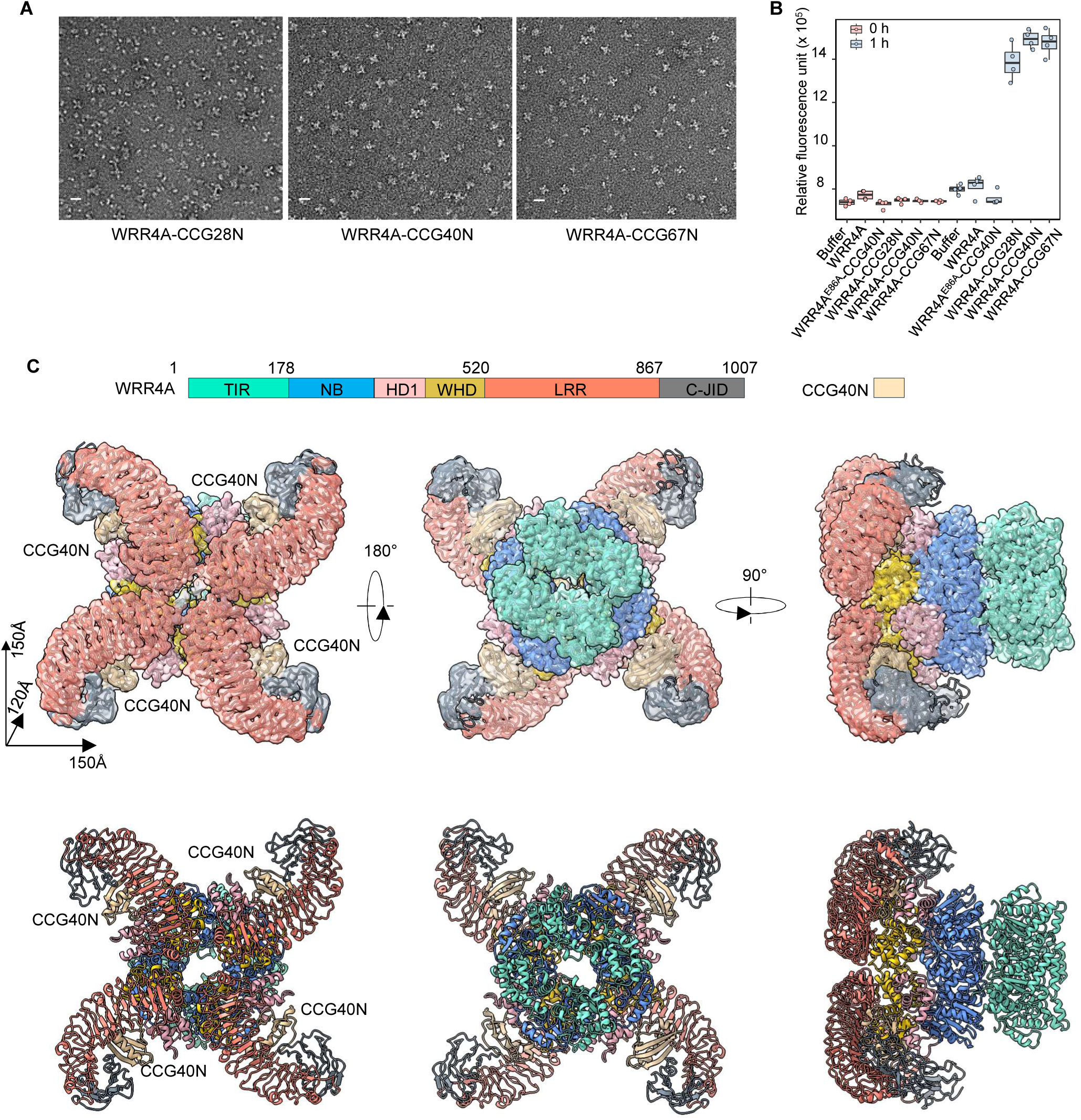
Cryo-EM analysis of the WRR4A-CCG40N resistosome. **(A)** Negative-stain electron microscopy of purified WRR4A complexes with CCG28N, CCG40N, and CCG67N (N-terminal recognition domains of CCG effectors). Cross-shaped particles characteristic of WRR4A oligomers are observed in all three samples, with varying degrees of particle homogeneity. Scale bars, 20 nm. **(B)** NADase activity of purified WRR4A complexes measured using the fluorescent NAD⁺ analog ε-NAD. Fluorescence was recorded immediately after substrate addition and after 1 h of incubation. WRR4A alone and the catalytic mutant WRR4A^E86A^ in complex with CCG40N (WRR4A^E86A^-CCG40N) served as negative controls. Data are from four replicates. **(C)** Cryo-EM reconstruction (top) and structural model (bottom) of the WRR4A-CCG40N resistosome complex. Individual WRR4A domains and the bound CCG40N effector are shown in distinct colours.

We performed cryo-EM structure determination of the WRR4A-CCG40N resistosome based on its overall sample quality. The tetrameric WRR4A-CCG40N resistosome, reconstructed with C2 symmetry (3.6 Å resolution, Fig. S2, Supplementary Table 2), displays ordered TIR-NLR domains with an overall arrangement closely resembling RPP1 and ROQ1 resistosome structures(Ma et al., 2020; Martin et al., 2020). The resistosome measures approximately 150 × 150 × 120 Å, with extensive oligomerization mediated by the TIR and NB-ARC domains (Fig. 2C). Side-chain-level density is resolved for most of WRR4A, except at the LRR periphery, including the C-JID domain, due to flexibility. The final model comprises residues 21-94 of the effector and residues 8-992 of WRR4A (1007 total residues), with side-chain details throughout most of the effector. Additional density at the NB domain was modelled as ADP, as ATP would likely create steric clashes at the γ-phosphate position (Fig. S3A). ADP binding at the NB domain has also been reported for RPP1, but not ROQ1(Ma et al., 2020; Martin et al., 2020), and correlates with an amino acid substitution in the P-loop motif from arginine (R297 in ZAR1 and R329 in ROQ1) to glutamate (E400 in RPP1), which abolishes γ-phosphate coordination. Consistent with this, WRR4A contains a glutamate (E325) at this position (Fig. S3B, C).

Like other TIR-NLRs(Ma et al., 2020; Martin et al., 2020), the four WRR4A protomers are brought together mainly through extensive interactions between their NB-ARC modules (Fig. 3A, E), with additional stabilization provided by TIR-TIR (Fig. 3B), LRR-LRR (Fig. 3F), and TIR-NB interdomain contacts (Fig. 3C). In addition to these interfaces that were well-characterized in previous studies, we observed well-resolved density for the loop connecting the TIR and NB domains (loop^TIR-NB^), which engages the NB domain of a neighbouring WRR4A protomer, forming an additional inter-protomer interaction (Fig. 3D). Disruption of this interface by a quadruple substitution within the loop^TIR-NB^ (WRR4A^N176A,T178A,R181A,D182A^) or a single mutation WRR4A^D182I^ abolished both effector-triggered WRR4A-mediated cell death (Fig. 3G) and WRR4A oligomerization (Fig. 3H), indicating a critical role for this loop in stabilizing the resistosome. Interestingly, the loop^TIR-NB^ is also resolved in the ADP-bound RPP1 tetramer but is absent in the ATP-bound ROQ1 tetramer (Fig. S3D), suggesting that this inter-protomer interaction may differ among TIR-NLRs and nucleotide binding states.

**Fig. 3.**
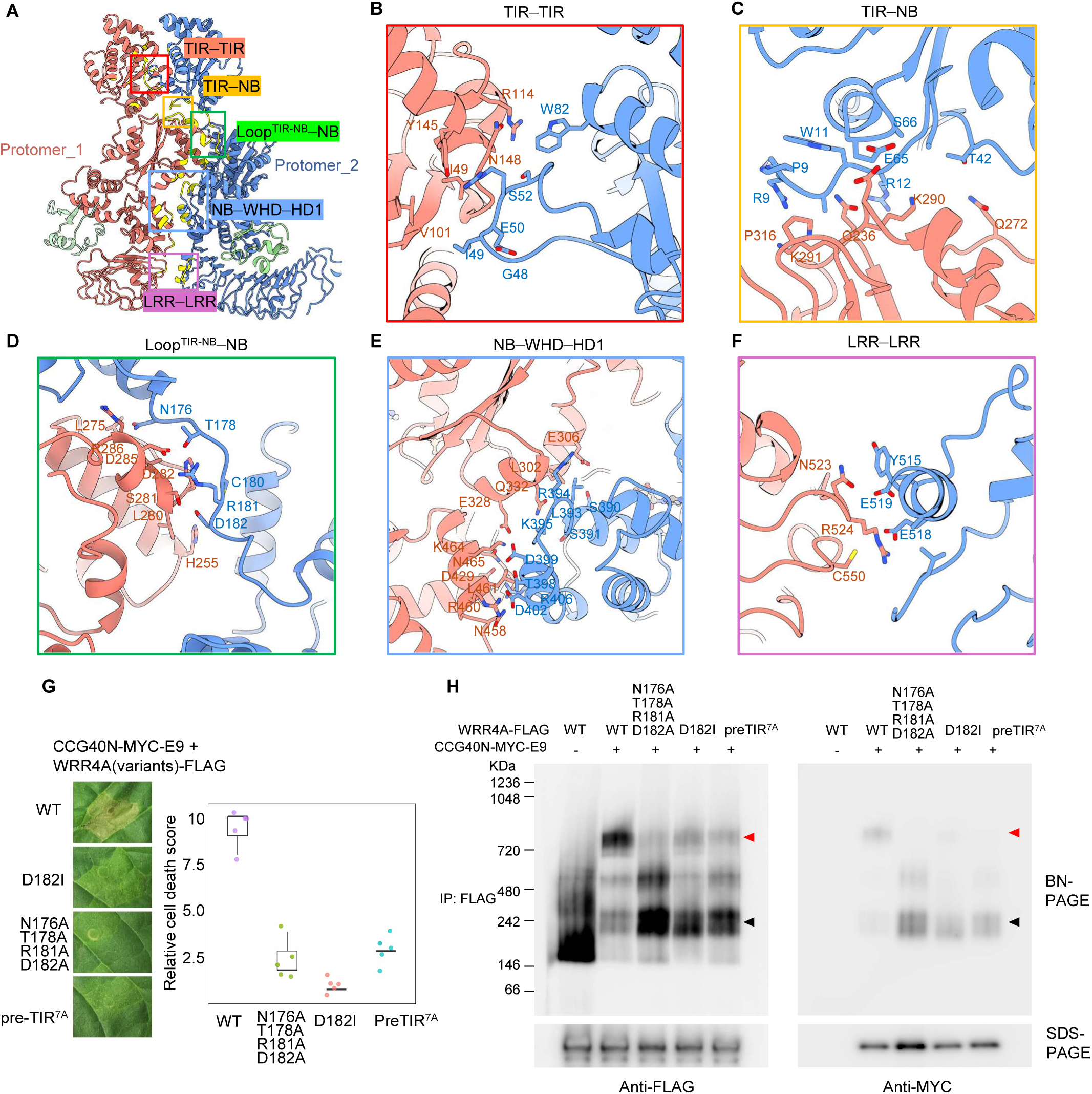
Assembly of tetrameric WRR4A resistosome. **(A)** Two adjacent protomers within the WRR4A-CCG40N resistosome, illustrating inter-protomer assembly. Interacting residues were identified in ChimeraX using a van der Waals overlap threshold of ≥ - 0.4 Å and are highlighted in yellow. Five major inter-protomer interfaces were identified: TIR–TIR, TIR–NB, loop^TIR-NB^–NB, NB–HD1–WHD, and LRR–LRR interactions. (**B–F**) Close-up views of the major inter-protomer interfaces shown in (A), highlighting key residues involved in resistosome assembly. **(G)** Cell death responses mediated by WRR4A interface mutants. Mutations at the preTIR–NB and loop^TIR-NB^–NB interfaces abolished effector-triggered cell death. Pre-TIR7^A^ denotes alanine substitution of seven residues in the loop immediately N-terminal to the TIR domain (P8 to N14). Cell death was assessed two days after Agrobacterium infiltration in *Nicotiana tabacum*. **(H)** BN-PAGE analysis revealed impaired oligomerization of WRR4A mutant variants. Red and black arrows indicate WRR4A-CCG40N tetramers and heterodimers, respectively. FLAG-tagged WRR4A variants were co-expressed with CCG40-MYC-E9 in *N. benthamiana eds1* mutants to prevent cell death.

In addition to the major interfaces described above, we observed density corresponding to an unstructured region immediately N-terminal to the TIR domain (residues P8-R9-N10-W11-R12-Y13-N14) that contacts the NB domain of a neighbouring protomer (Fig. 3C; Fig. S3E, F). This segment follows a tandem N-terminal serine repeat whose pattern is conserved across multiple TIR-NLRs (Fig. S3E). Although residues N-terminal to R12 could not be confidently modelled, their proximity to the inter-protomer interface suggests a contribution to oligomer stabilization (Fig. S3F). Consistent with this interpretation, alanine substitution of residues P8-N14 (PRNWRYN→AAAAAAA; pre-TIR^7A^) abolished effector-triggered WRR4A oligomerization and the associated cell death response (Fig. 3G, H), demonstrating that this region contributes to resistosome stability. By contrast, mutation of the tandem serine residues (WRR4A^S3-7A^) did not impair WRR4A-mediated cell death (Fig. S3G). The conservation of this pre-TIR region suggests a common mechanism contributing to TIR-NLR oligomerization. Consistent with these observations, AlphaFold3(Abramson et al., 2024) modelling of the WRR4A^TIR-NB-ARC^ tetramer suggests inter-protomer contacts involving the loop^TIR-NB^ and pre-TIR regions with high predicted confidence (Fig. S3H, I).

The TIR domains in the WRR4A resistosome were well resolved and adopt a dimer-of-heterodimers arrangement (Fig. S3J), consistent with that of RPP1 and ROQ1(Ma et al., 2020; Martin et al., 2020). In the reported RPP1 resistosome structure, ATP supplied during purification occupies the NAD⁺-binding sites within the TIR domains. In contrast, no nucleotide density was observed at these sites in the WRR4A structure (Fig. S3K), likely because no exogenous ATP or NAD⁺ was supplied during purification and any endogenous substrate had been catalysed.

### Molecular basis of CCG40N recognition by WRR4A

Consistent with structural predictions obtained using AlphaFold2, CCG40N (residues 20-130) in the cryo-EM structure exhibits a ferredoxin-like fold comprising an N-terminal four-stranded antiparallel β-sheet packed against a long α-helix, forming the rigid core of the effector. The C-terminus consists of a loop containing a short helix connected to the ordered domain (Fig. 4A). In the cryo-EM reconstruction, the helix and β-sheet regions of the effector are resolved to side-chain level. The ellipsoidal ferredoxin-like domain of CCG40 occupies the inner concave surface of the WRR4A LRR domain (Fig. S4A). WRR4A engages CCG40 through three domains – the WHD, LRR, and C-JID – with a total buried surface area of 2530 Å² upon complex formation (Fig. 4A, Fig. S4A) and co-IP data suggest that the LRR and C-JID are the primary interacting regions (Fig. S4B).

**Fig. 4.**
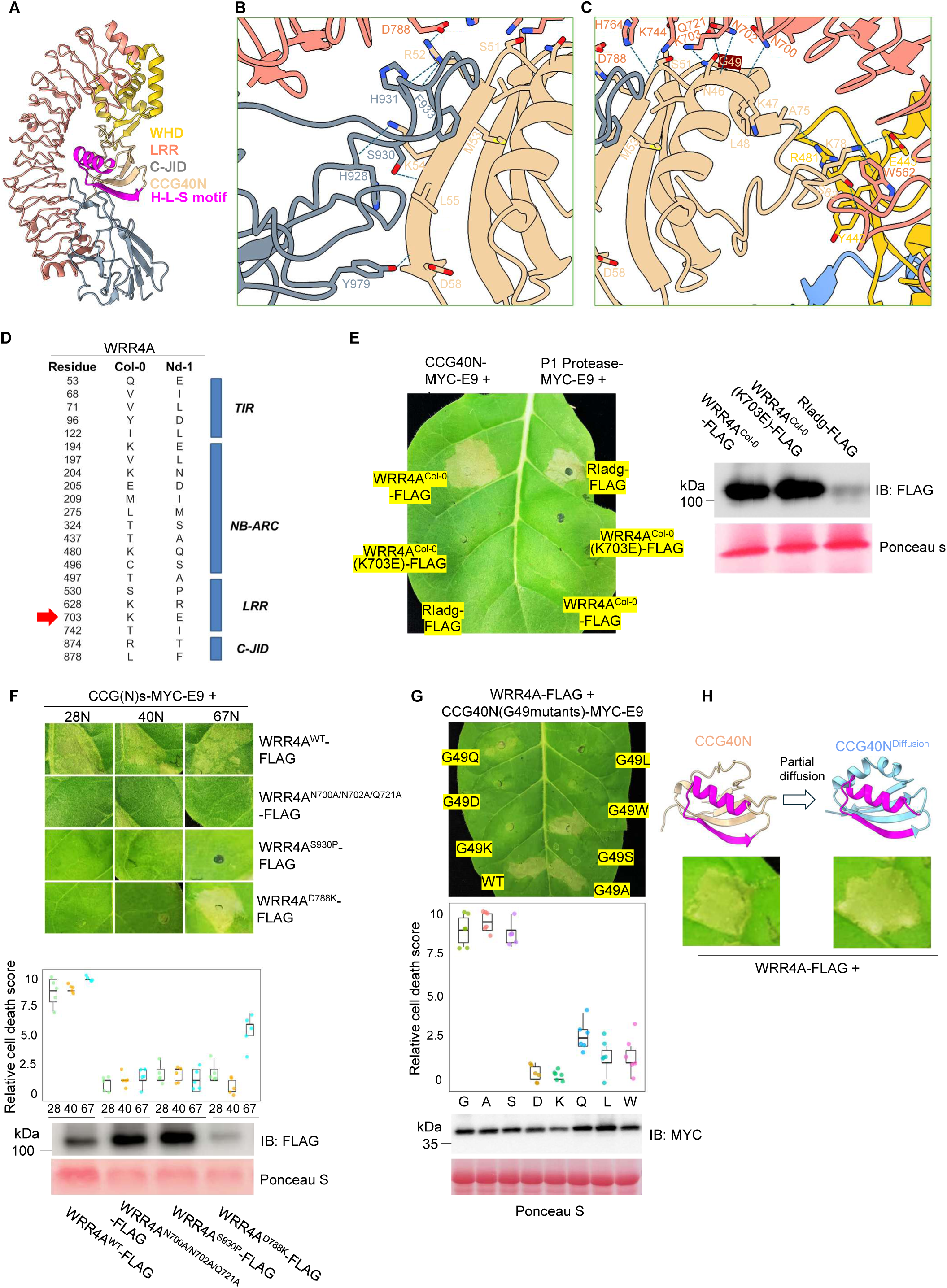
WRR4A-mediated recognition of CCG40N. **(A)** Structural model highlighting the major interaction interface between WRR4A and CCG40N. WRR4A engages CCG40N primarily through its LRR and C-JID domains, contacting the α1-loop-β2 region (helix-loop-strand [H-L-S] motif) of CCG40N (magenta). **(B, C)** Close-up views of specific interactions at the WRR4A-CCG40N interface. **(D)** Sequence polymorphisms between WRR4A^Nd-1^ and WRR4A^Col-0^. The K703E substitution at the WRR4A-CCG40 interaction interface is highlighted. **(E)** The K703E substitution in WRR4A^Col-0^ abolishes recognition of CCG40N. FLAG-tagged WRR4A variants were co-expressed with CCG40-MYC-E9 in *N. tabacum*. Cell death was assessed two days after infiltration. Protein expression was verified in a separate experiment in which WRR4A variants were expressed in *N. benthamiana eds1* mutants to avoid cell death. An unrelated potato NLR RIadg, and its cognate effector P1 protease are used as negative controls. **(F)** Mutation analysis of WRR4A interface residues confirms their contribution to CCG recognition. FLAG-tagged WRR4A variants were individually co-expressed with CCG28N-, CCG40N-, or CCG67N-MYC-E9, and cell death was assessed two days post infiltration. Protein expression was confirmed by immunoblotting from separately infiltrated *N. benthamiana eds1* mutants to prevent cell death. Data points are from five individual leaves. **(G)** Cell death assays assessing the contribution of the conserved “G” within the CCG motif to WRR4A-mediated recognition. Cell death was scored two days post infiltration. Data points are from five individual leaves. Protein expression was confirmed using separate infiltrations in *N. benthamiana eds1* mutants to avoid cell death. **(H)** Artificially designed CCG40N^Diffusion^ can be recognized by WRR4A. CCG40N^Diffusion^ was designed using partial diffusion with RFdiffusion, generating a ferredoxin-like protein with a randomized sequence while retaining the H-L-S motif of CCG40N (Magenta). AlphaFold confirmed the intended fold (left). Cell death was assessed two days after co-infiltration with WRR4A-FLAG (right).

Using a 4.5 Å distance cutoff between the effector and WRR4A to define non-covalent contacts, we identified interactions between 31 residues of WRR4A and 23 residues of the effector (Fig. 4B, C; Supplementary Table 1). In CCG40N, residues 37-47 form a long α-helix (α1), which is connected to the β2 strand (residues 52-58) via an intervening loop; together, these elements constitute a structural element we refer to as the Helix-Loop-Strand (H-L-S) element (Fig. 4A). This element forms the principal interface for WRR4A recognition. Notably, the β2 strand (residues 52-58) is oriented such that its backbone atoms are presented toward the loops of the WRR4A C-JID domain in the complex. As a result, much of the molecular recognition is potentially mediated by contacts between WRR4A LRR and C-JID domains and backbone atoms of the CCG40N within residues 46-58 (N46, G49, A50, S51, M53, L55, and D58) (Fig. 4B, C). Sequence analysis reveals substantial divergence in this region among CCGs (Fig. S4C), indicating that backbone-mediated recognition at multiple positions enables WRR4A to tolerate sequence variation and underlies its broad recognition capacity. In addition, residues including K703 and D788 within the WRR4A LRR domain form side chain-side chain interactions with the H-L-S element of CCG40N. Together, these observations indicate that WRR4A recognizes CCG40N by a combination of its shape (backbone atoms at defined positions), and side chains of surface residues.

The structure information explains why WRR4A^Nd-1^, a natural WRR4A allele, is non-functional. WRR4A was originally identified using an F2 population derived from a cross between the Col-0 (*A. candida* resistant) and Nd-1 (*A. candida* susceptible), indicating that WRR4A^Nd-1^ lacks function(Borhan et al., 2010). Our previous study showed that WRR4A^Nd-1^ fails to bind and recognize CCG effectors detected by WRR4A^Col-0^. Replacing the WRR4A^Nd-1^ LRR domain with that from WRR4A^Col-0^ restores both CCG interaction and recognition, indicating that the defect lies in effector recognition rather than in oligomerization or TIR-domain activity(Redkar et al., 2023). Comparison of interface residues within LRR domains of the two alleles revealed that K703 in WRR4A^Col-0^ is substituted with glutamate in WRR4A^Nd-1^ (Fig. 4D). Introducing K703E into WRR4A^Col-0^ abolished CCG recognition without destabilizing the protein (Fig. 4E), demonstrating that the substitution accounts for the loss of function in WRR4A^Nd-1^, validating the structural model.

To assess the functional importance of backbone-mediated recognition, we introduced a triple substitution in the LRR domain (N700A/N702A/Q721A) designed to disrupt side chain-backbone interactions at the WRR4A-CCG40N interface (Fig. 4C). This mutation abolished WRR4A-mediated recognition of CCG28N, CCG40N, and CCG67N, highlighting the requirement for these contacts in effector recognition (Fig. 4F). In parallel, an S930P substitution was introduced in the C-JID region that interacts with the backbone atoms of β2 strand from the H-L-S element; the substitution removes a backbone hydrogen-bond donor and restricts local geometry, and this mutation abolished recognition of all three effectors (Fig. 4F).

In addition to backbone-mediated contacts, side chain-side chain interactions further contribute to WRR4A recognition and provide specificity. D788 in the LRR domain forms a charged interaction with CCG40N R52 (Fig. 4B), a residue conserved among several CCG effectors (Fig. S4C). D788K abolished WRR4A-mediated recognition of CCG28N and CCG40N, but not CCG67N, which lacks this conserved arginine (Fig. 4F, Fig. S4C).

On the effector side, residue GLY49 of CCG40N drew particular attention as the conserved “G” within the CCG motif (Fig. S4C). Mutation of this conserved GLY in CCG28 does not abolish WRR4A-mediated recognition(Redkar et al., 2023). In the WRR4A-CCG40N structure, GLY49 of CCG40N lies adjacent to the WRR4A LRR domain and engages in backbone-mediated contacts with the side chain of Q721 (Fig. 4C, Fig. S4D). Consistent with a backbone interaction, CCG40^G49S^ and CCG40^G49A^ remained recognizable by WRR4A, whereas substitutions with bulkier or charged residues at this position abolished recognition (Fig. 4G). These results indicate that spatial constraints at this interface, rather than the glycine residue per se, are key determinants of recognition, with small side chains compatible with WRR4A engagement.

To further validate the importance of the H-L-S element of CCG40N in recognition, we used the partial diffusion function of RFdiffusion(Watson et al., 2023) to generate a ferredoxin-fold with randomized sequence but still bearing the H-L-S element of CCG40N, creating an artificial effector, CCG40N^Diffusion^ (Fig. 4H; Fig. S4E). Structurally similar to CCG40N but sharing sequence only in the H-L-S region (Fig. S4E), CCG40N^Diffusion^ triggered robust HR when co-expressed with WRR4A in *N. tabacum* (Fig. 4H). These results further demonstrated that WRR4A recognizes the H-L-S element of a ferredoxin-like fold.

### Cryo-EM structure of WRR4A-resistosome with a weakly bound effector (CCG28N)

To elucidate how different CCGs are recognized, we determined a second WRR4A resistosome structure in complex with CCG28N. CCG28N was selected over CCG67N because the WRR4A-CCG28N protein sample exhibited distinct characteristics in negative-stain EM analysis (Fig. 2A). Surprisingly, cryo-EM reconstruction of the WRR4A-CCG28 complex revealed that the effector binding site is largely empty with only faint, fragmented density in 3D-reconstruction, while the complete WRR4A resistosome is resolved to side-chain level, resembling an apo-resistosome state (Fig. 5A, Fig. S2, Supplementary Table 2). This observation indicates that, unlike the stably bound CCG40N, CCG28N binds weakly to a subset of WRR4A protomers with insufficient occupancy to generate coherent density. To generate a structure model that represent the complete complex, we performed rigid-body placement of the AlphaFold2-predicted CCG28N model into the WRR4A apo-resistosome map. At adjusted map thresholds (0.0025), weak density corresponding to the overall shape of the effector was visible, facilitating its rigid-body placement (Fig. S5A, Fig. 5B). Superposition with WRR4A-CCG40N structure revealed that the binding orientation and pose of CCG28N and CCG40N are similar, suggesting that WRR4A recognizes two CCGs with similar mechanisms (Fig. 5C). Consistent with WRR4A-CCG40N, the WRR4A protomers likely bind ADP at their NB domain in the WRR4A-CCG28N structure (Fig. S3A).

**Fig. 5.**
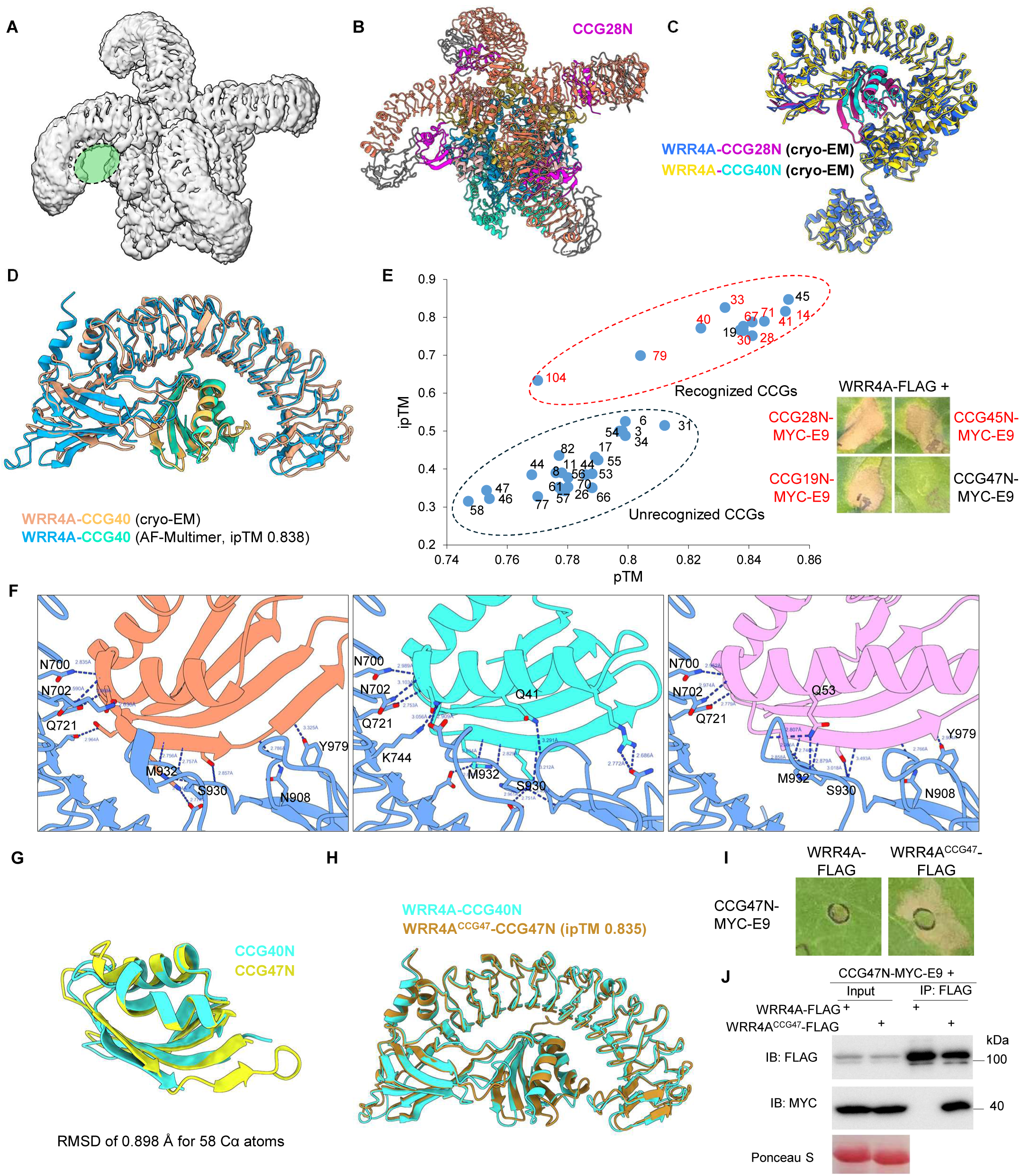
Structural basis of WRR4A-mediated broad recognition. **(A)** Cryo-EM density of WRR4A-CCG28N complex showing lack of density in the effector binding site. **(B)** Model of the CCG28N-bound WRR4A resistosome. The highest ranked AlphaFold2 model of CCG28N was rigid-body fitted into the cryo-EM map of the WRR4A apo-resistosome using an adjusted density threshold (0.0025), at which weak but discernible effector density was visible. **(C)** Superposition of WRR4A-CCG protomers from the CCG28N- and CCG40N-bound resistosomes reveals highly similar effector-binding modes. **(D)** The AlphaFold-Multimer model of WRR4A-CCG40N closely matches its cryo-EM structure. LRR and C-JID domains of WRR4A were used for predicting the interaction with CCG40N. **(E)** AlphaFold-Multimer accurately discriminates between recognized and unrecognized CCG effectors. Previously characterized CCGs were modelled using their N-terminal domains together with the LRR and C-JID region of WRR4A. The recognition of CCG19N and CCG45N by WRR4A were re-examined, which were previously classified as unrecognized by WRR4A yet yielded high predicted ipTM scores. Cell death was evaluated two days post co-infiltration of CCG effectors with WRR4A. **(F)** Comparison of AlphaFold-Multimer models of WRR4A bound to CCG28N, CCG40N, and CCG67N reveals conserved backbone hydrogen-bonding patterns at the interaction interface. **(G)** CCG47N, which is not recognized by WRR4A, adopts a structure highly similar to that of CCG40N. **(H)** AlphaFold-Multimer prediction of the engineered WRR4A^CCG47^ variant with CCG47N. **(I)** WRR4A^CCG47^ recognizes CCG47N in *N. tabacum*. Cell death was assessed two days post infiltration. Results are representative of five biological replicates. **(J)** Co-immunoprecipitation assays demonstrate interaction between WRR4A^CCG47^ and CCG47N. *N. benthamiana eds1* mutant plants were infiltrated to avoid cell death. Tissue was harvested three days post infiltration, and total proteins were immunoprecipitated with anti-FLAG beads and analyzed by immunoblotting with anti-FLAG and anti-MYC antibodies.

Given that CCG28N triggers robust cell death comparable to CCG40N (Fig. 1D, E), and that the WRR4A-CCG28N complex exhibits similar enzymatic activity to WRR4A-CCG40N (Fig. 2B), resistosome formation appears unaffected by weaker CCG28N binding. The substantial fraction of effector-free WRR4A tetramers (apo-resistosomes) suggests that sustained effector binding is not required to maintain resistosome integrity. Instead, the activated WRR4A tetramer appears self-stabilized through inter-protomer interactions, consistent with auto-active NLR mutants that oligomerize without effector binding(Liu et al., 2024; Guo et al., 2025).

### The structural basis of WRR4A-mediated broad recognition

Due to the limited resolution of CCG28N in the cryo-EM structure, we used AlphaFold-Multimer(Evans, 2021) to predict how WRR4A interactions with different CCGs. Only the LRR and C-JID domains of WRR4A and the N-terminal regions of CCGs were used, as these mediate the primary contacts. Remarkably, AlphaFold-Multimer produced high-confidence models of the WRR4A-CCG40N complex (ipTM = 0.838) that closely matched the cryo-EM structure (Fig. 5D). To further evaluate the accuracy of AlphaFold-Multimer predictions, we extended our predictions to additional CCGs whose recognition by WRR4A had been experimentally validated in a previous study(Redkar et al., 2023). Notably, AlphaFold-Multimer accurately distinguished between recognized and unrecognized CCGs, with recognized CCGs clustering together with high ipTM scores and unrecognized CCGs clustering together with low ipTM scores (Fig. 5E).

During this *in silico* interaction screening, we identified two CCGs, CCG19 and CCG45, with high ipTM scores despite being previously characterized as unrecognized by WRR4A(Redkar et al., 2023) (Fig. 5E). To resolve this discrepancy, we re-cloned and co-expressed them with WRR4A, and both triggered a strong HR, suggesting the earlier results were false negatives. Furthermore, two additional WRR4A-recognized CCGs (CCG14 and CCG41) were identified by proximity-labelling coupled with mass spectrometry (PL-MS) using WRR4A-TurboID transgenic Arabidopsis *eds1* mutants infected with *A. candida* isolate Ac2V, followed by biotin labelling (Fig. S5C, D)(Szymansky, 2025). AlphaFold-Multimer predicted high-confidence interactions for both effectors (Fig. 5E), further validating its predictive accuracy and expanding the repertoire of WRR4A-recognized CCGs.

In the WRR4A-CCG40 cryo-EM structure, WRR4A engages the backbone atoms of CCG40N at multiple sites, likely allowing tolerance of residue substitutions and enabling broad recognition. Leveraging the accuracy of AlphaFold-Multimer predictions, we tested this hypothesis by comparing predicted models of WRR4A complexed with CCG28N, CCG40N, and CCG67N. Consistent with our two cryo-EM structures, all three effectors adopt similar binding modes, with the H-L-S elements in close contact with the LRR and C-JID domains of WRR4A (Fig. 5F). We next examined hydrogen bonds between WRR4A and the H-L-S elements of three CCGs, which are critical for binding specificity. Despite sequence variation, the predicted contacting regions within different CCGs are similar. Notably, inter-molecule hydrogen bonds are predicted to occur at nearly identical positions across the three complexes (Fig. 5F). Closer inspection revealed that most bonds involve backbone atoms of the effectors and sidechain or backbone atoms of residues on the receptor, especially between CCG67N and WRR4A. Specifically, within the central LRR region, residues N700/N702/Q721 of WRR4A, which were proven critical for recognition (Fig. 4G), form hydrogen bonds with backbone atoms of the effectors. Consistent with the WRR4A-CCG40N cryo-EM structure, a second set of backbone-mediated interactions occurs between the effector β2 strand and the WRR4A C-JID domain. The β2 strand presents its backbone atoms toward C-JID loops, forming a ladder of backbone hydrogen bonds (Fig. 5F).

Together, we propose that WRR4A’s broad-spectrum recognition arises from extensive backbone-mediated interactions with multiple defined positions within the ferredoxin-like fold of CCG effectors, enabling recognition based primarily on conserved structural features rather than sequence homology. Additional sidechain and electrostatic interactions likely modulate binding affinity and confer effector-specific discrimination. Despite recognizing over ten CCGs, WRR4A retains specificity: most CCG effectors are still not recognized (Fig. 5E), and host ferredoxin-like proteins are presumably excluded to prevent autoactivation. This specificity may arise from a combination of factors, including side-chain incompatibilities and steric hindrance at closely contacting regions that impair close contact (such as aromatic amino acid replacements at the conserved “G” position shown in Fig. 4G). WRR4A balances broad structural recognition with precise discriminatory mechanisms that ensure effective defense while avoiding self-targeting.

### Structure-guided engineering enables WRR4A to gain new recognition

With structural information and the aid of accurate AlphaFold-Multimer predictions, we explored the possibility of engineering WRR4A to acquire novel recognition specificity. CCG47 is a CCG effector that is not recognized by WRR4A (Fig. 5E). Despite low sequence identity to CCG40 (21%; Fig. 1A), the predicted CCG47N structure closely resembles that of CCG40N (RMSD = 0.898 Å for 58 Cα atoms; Fig. 5G). Consistent with the lack of recognition, AlphaFold-Multimer predicted an incorrect binding orientation for WRR4A-CCG47N with a low ipTM score (Fig. S5E).

Effectors can evade recognition by altering key recognition residues or by introducing steric hindrances that obstruct interactions. Given that WRR4A tolerates substantial sequence variation at the interaction interface, we reason that steric hindrance is likely more effective in disrupting WRR4A recognition. Comparison of interface residues between CCG40N and CCG47N revealed that ASN35 in CCG40N is replaced by an ASP in CCG47N. To potentially create a charge interaction with this ASP, the closely contacting GLU903 in WRR4A was mutated to LYS (Fig. S5F). Examination of the H-L-S element of CCG47N further revealed a tyrosine residue within the CCG motif positioned near the backbone-recognition region (corresponds to ASN46 in CCG40N) that might affect the interaction (Fig. S5F, G). Combined with E903K, we performed an *in silico* mutation screening of surface-exposed residues on the WRR4A LRR domain in close contact with this tyrosine (Fig. S5F), using Alphafold-Multimer ipTM scores as a readout. This analysis identified K744A as a favourable substitution which, in combination with E903K, increased the WRR4A-CCG47N ipTM score from 0.35 to 0.6. Finally, a WRR4A^D981N^ substitution further increased the ipTM score. The original ASP981 interacts with ARG34 in CCG40N, whereas the corresponding position in CCG47N is a GLU, the D981N substitution might promote alternative interactions. A WRR4A variant bearing all three substitutions (WRR4A^K744A/E903K/D981N^) exhibited a high ipTM score of 0.835 with CCG47N, with CCG47N adopting a similar binding mode to CCG40N in the prediction (Fig. 5H). HR assays confirmed that CCG47N is recognized by this engineered variant, designated WRR4A^CCG47^ (Fig. 5I). co-IP further demonstrated the interaction between WRR4A^CCG47^ and CCG47N (Fig. 5J). Notably, WRR4A^CCG47^ retained recognition of strongly interacting effectors (CCG28N, CCG30N, CCG40N, and CCG67N) but lost recognition of weakly interacting CCG33N and CCG71N (Fig. S5H).

Together, these results demonstrate that structure-guided engineering informed by AlphaFold predictions can successfully reprogram NLR recognition specificity. In contrast to previous approaches that relied on a combination of structural insights and natural genetic variation(Forderer et al., 2022; Lawson et al., 2025), the strategy depends exclusively on structural information, making it broadly applicable for engineering novel NLR specificities even in the absence of prior genetic or evolutionary data.

### Cryo-EM data support an AlphaFold prediction model for WRR4A resting state

While resting (pre-activation) state structures of CC-NLRs before effector recognition have been experimentally resolved (ZAR1 and NRC2)(Wang et al., 2019b; Ma et al., 2024; Selvaraj et al., 2024), no experimental structure of a resting state TIR-NLR is currently available, hindering the understanding of their activation process upon effector binding. TIR-NLRs like WRR4A may adopt distinct resting state conformations from CC-NLRs due to their TIR domains and additional C-JID domains. Blue-native PAGE analysis suggests that WRR4A exists predominantly as a monomer prior to activation (Fig. 1G; Fig. 3H). Negative staining analysis revealed WRR4A resting state as small particles that likely represent monomers (Fig. S6A). We sought to determine the resting state structure using cryo-EM. Although the resting state WRR4A could be purified to homogeneity using a tandem purification strategy (Fig. S1C), its relatively small size (∼100 kDa), together with strong preferred orientation, precluded high-resolution cryo-EM structure determination in our analysis. Nevertheless, 2D class averages of cryo-EM images of resting WRR4A revealed well-defined monomeric particles with secondary structure details, including β-strand separation in the LRR domain. Individual domains could be distinguished apart from the TIR domain, for which no clear density was observed (Fig. 6A, Fig. S6B).

**Fig. 6.**
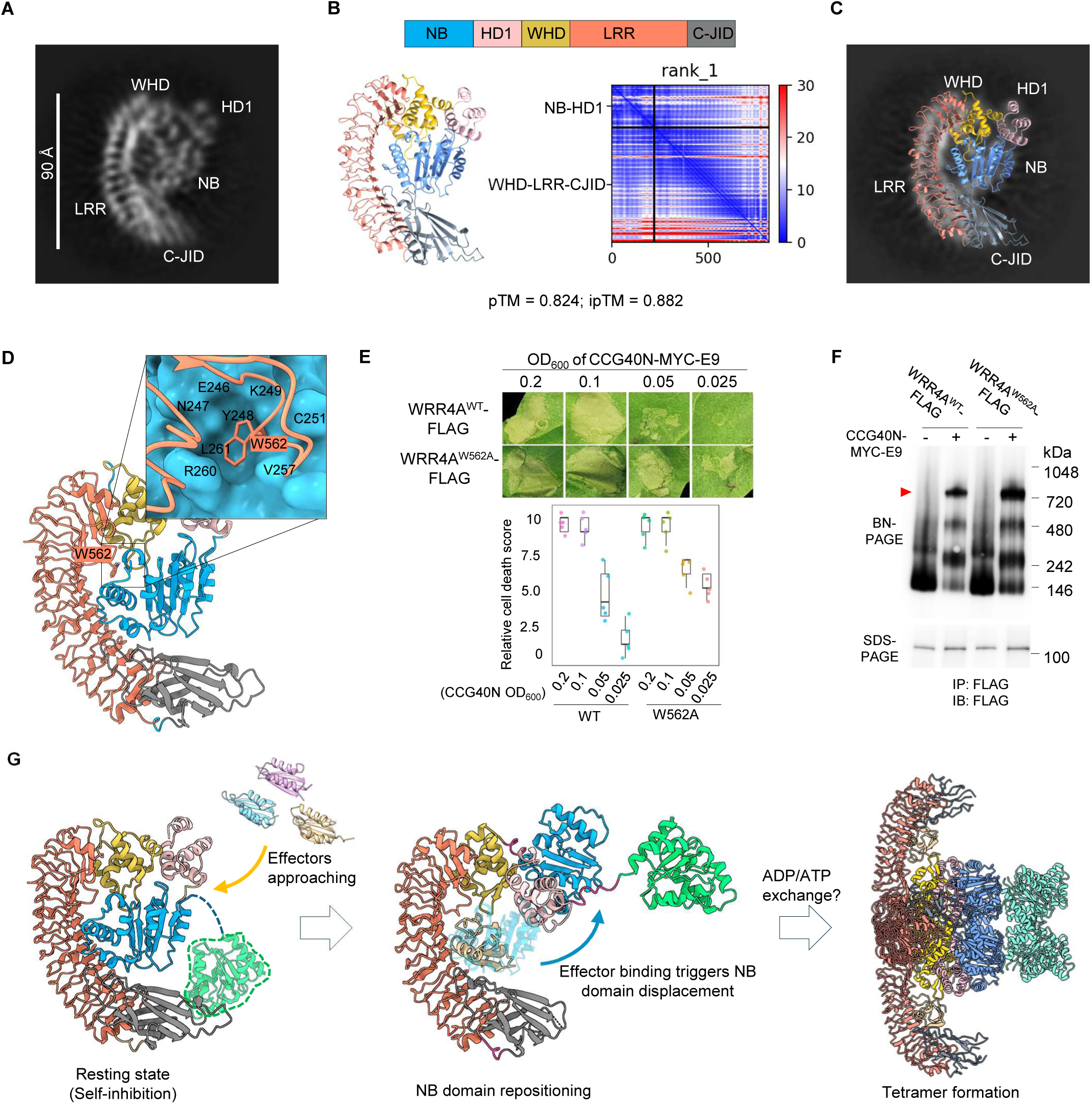
A plausible model for the WRR4A resting state and activation mechanism. **(A)** Representative cryo-EM 2D class averages of resting-state WRR4A reveal monomeric particles with discernible secondary structure features. All domains are distinguishable except the TIR domain, for which no clear density is observed. **(B)** AlphaFold-Multimer predicted model of the WRR4A monomer closely matches the experimental resting state 2D class averages. The TIR domain was not included in the prediction. **(C)** Alignment of the AlphaFold-predicted WRR4A monomer model with 2D class average projections of resting-state WRR4A. **(D)** In the predicted resting-state model, W562 in an extended loop between LRR2-LRR3 inserts into a pocket on the NB domain, stabilizing the NB–LRR interaction in the resting state. **(E)** Cell death assays show that the W562A mutation sensitizes WRR4A. Agrobacterium expressing WRR4A-FLAG were co-infiltrated with serial dilutions of strains expressing CCG40N-MYC-E9. WRR4A^W562A^ remains responsive at OD_600_ = 0.05 of CCG40N. Agrobacterium carrying WRR4A variants have a consistent infiltration OD_600_ = 0.2. Cell death was scored two days post infiltration. **(F)** Blue-native PAGE analysis shows enhanced tetramer formation by WRR4A^W562A^ compared with wild-type WRR4A at low effector concentrations. The red arrow indicates the WRR4A-CCG40N tetrameric complexes. FLAG-tagged WRR4A variants (OD_600_ = 0.2) were co-infiltrated with CCG40N-MYC-E9 (OD_600_ = 0.05) in *N. benthamiana eds1* mutants. Results are representative of three independent experiments. **(G)** Proposed model for WRR4A activation. In the resting state, the NB domain is held within the concave surface of the LRR domain, overlapping with the effector-binding site through interactions with the LRR and C-JID domains. Binding of CCG effectors competitively displaces the NB domain, promoting nucleotide exchange, conformational rearrangement, and irreversible resistosome assembly.

Our attempts to predict full-length WRR4A resting states using AlphaFold2 or AlphaFold3 repeatedly yielded activated protomer conformations that were not consistent with our 2D-class averages. We reasoned that full-length input might constrain conformational sampling and bias predictions toward activated states. Since HD1 and WHD likely undergo major rearrangements upon activation(Wang et al., 2019a; Wang et al., 2019b), we divided WRR4A at the loop linking the HD1 and WHD domains, creating two chains: TIR-NB-HD1 (or NB-HD1) and WHD-LRR, to allow more flexible positioning during structure prediction. This resulted in high-confidence conformations that were more compact and distinct from the activated state (Fig. 6B, Fig. S6C). Inclusion of the TIR domain reduced the prediction confidence, with the PAE plot indicating poor positional confidence for the TIR domain relative to the other domains (Fig. S6C), consistent with the absence of TIR density in the cryo-EM 2D class averages. We infer the TIR domain remains conformationally flexible in the resting state. Several lines of evidence support that the predicted model (excluding the TIR domain) represents a *bona fide* resting state conformation. First, projection matching showed close agreement between the model and experimental 2D class averages (Fig. 6C). A low-resolution 3D reconstruction from these particles resolved individual domains and suggested that the TIR domain is disordered in the resting state, although precise domain placement was limited by the absence of resolved secondary-structure features (Fig. S6E). Second, superposition of the NB-ARC domain onto the resting state ZAR1 structure revealed a highly similar relative domain arrangement, consistent with an inactive conformation (Fig. S6F). Third, the conserved MHD motif was modelled in a configuration compatible with ADP binding, with the histidine positioned to accommodate the β-phosphate (Fig. S6G), a structure feature of NLR resting state(Bonardi et al., 2012).

In the predicted model of resting state, the NB domain of WRR4A nestles within the concave curvature of the LRR domain. This resting conformation is further stabilized by interactions with the C-JID domain and the WHD domain from two opposite sides. Specifically, two large loops of the C-JID domain interact with the NB domain in the resting state (Fig. 6B, Fig. S6D). Comparison of the WRR4A resting state with those of CC-NLRs (ZAR1 and NRC2)(Wang et al., 2019b; Ma et al., 2024; Selvaraj et al., 2024) reveals that the WRR4A^LRR^ domain adopts a more vertical orientation, lying roughly in the plane of the NB-HD1-WHD domains, in contrast to the LRR domains of ZAR1 and NRC2. This vertical orientation of the WRR4A LRR domain creates the spatial accommodation required for the C-JID to approach and engage the NB domain from above, thereby stabilizing the resting conformation - a configuration that would otherwise not be feasible (Fig. S6H). It is reasonable to speculate that other TIR-NLRs with a C-JID or NLRs with C-terminal integrated domains adopt resting states similar to WRR4A, allowing the integrated domain to approach and stabilize the NB domain until cognate effector binding occurs.

In the resting state, the NB domain is maintained in a constrained position through interactions with the LRR and C-JID domains, while the TIR domain remains conformationally flexible. Notably, the NB domain position in the resting state overlaps with the effector binding site in the activated WRR4A protomer. This suggests that competitive binding of CCG effectors displaces the NB domain, enabling nucleotide exchange and HD1 domain release, analogous to the ZAR1 activation mechanism (Fig. 6G)(Wang et al., 2019a; Wang et al., 2019b). In the predicted resting state structure, a tryptophan residue (W562) in an extended loop between LRR2-LRR3 inserts into a pocket on the NB domain, stabilizing the NB-LRR interface (Fig. 6D). Mutation to alanine did not trigger autoactivation but sensitized WRR4A, enabling activation at lower effector concentrations (Fig. 6E, F). These findings underscore the role of W562 in maintaining the resting state and demonstrate the potential to engineer hypersensitive NLR variants that respond more efficiently during early pathogen detection.

## DISCUSSION

Rapidly evolving pathogens can quickly overcome resistance conferred by a single resistance gene. However, WRR4A, an NLR with exceptional broad-recognition capacity, can provide both broad-spectrum and potentially durable resistance to *Albugo candida*(Borhan et al., 2010; Redkar et al., 2023). We investigated the structural basis of this broad recognition capacity, revealing a shape recognition mechanism not previously seen in other NLRs. In addition, we defined a plausible monomeric resting state conformation of WRR4A, representing a TIR-NLR resting state with a C-JID domain and revealing certain key differences from reported CC-NLR resting states. This resting state structure facilitates understanding of how TIR-NLRs with C-JID motifs maintain their resting forms and are activated by effector binding. Furthermore, these structural insights enabled engineering of two WRR4A variants—one with altered recognition specificity and one with enhanced sensitivity to low effector concentrations—demonstrating the potential of structure-informed resistance engineering.

### Structural basis for broad range effector recognition by WRR4A

From analysis of the cryo-EM structures and *in silico* predictions of CCG binding to WRR4A, we propose a three-step recognition mechanism. First, the receptor imposes a size constraint, selecting proteins that fit within the space defined by the C-JID, LRR, and WHD domains. Second, through interactions with backbone atoms in the H-L-S element of the CCG effector, it selects for the ferredoxin-like fold. Third, recognition of surface side chains enables specific selection of pathogen effectors while excluding host proteins with this fold. This mode of recognition provides a structural explanation for how a single NLR can achieve unusual broad effector specificity while retaining selectivity. Oomycete and fungal pathogens frequently secrete effectors that are structurally conserved yet sequence-divergent(Seong and Krasileva, 2021, 2023; Lawson et al., 2025). These effectors often adopt stable protein scaffolds (*e.g.* ferredoxin-like fold, RNase-like fold) that tolerate substantial surface variation, enabling the evolution of diverse virulence functions and facilitating immune evasion. Our structural analysis indicates that instead of relying on sequence-specific side-chain interactions that are prone to mutational escape, WRR4A recognizes a conserved structural signature of CCG effectors by interactions with the effector backbone, enabling recognition of sequence-diverged effectors. These findings extend current models of NLR specificity and suggest that fold-based recognition can contribute to broad and potentially durable plant immune responses.

Backbone recognition is employed by many enzymes to achieve broad substrate specificity. For instance, peptidyl-tRNA hydrolase (PTH) forms sidechain-backbone hydrogen bonds with its peptide substrates to accommodate peptides of diverse sequences(Schmitt et al., 1997). Glycosyltransferases also form sidechain-backbone hydrogen bonds with their substrates through a series of asparagine and glutamine residues (Asn/Gln ladder) in their tetratricopeptide repeats (TPRs)(Levine et al., 2018; Zhu et al., 2022). A short Asn/Gln ladder (N700/N702/Q721) is also present in WRR4A and mediates backbone interactions with CCGs. These residues are favoured in backbone interactions because their side chains can act as both hydrogen bond donors and acceptors to interact with backbone carbonyl oxygens or amide hydrogens.

The conservation of the N-terminal ferredoxin-like fold among CCG effectors suggests an important functional role for these effectors. One possibility is that this fold serves as a translocation signal required for effector delivery into host cells, whereas the more divergent C-terminal domains mediate effector-specific virulence functions. Recognition of this conserved fold, rather than the variable remaining domains, enables the broad recognition capacity of WRR4A.

### Recognition of full-length CCGs

We performed most experiments using the ∼100-amino-acid N-terminal recognized domain of CCGs. Previous studies demonstrated that the remaining domains of CCGs are not required for recognition(Redkar et al., 2023). In AlphaFold2 models of full-length CCGs, the recognized N-terminal ferredoxin-like domain is connected to the C-terminal region by a long, flexible linker and remains fully solvent-exposed (Fig. 1C), suggesting that full-length CCGs should interact with WRR4A in a manner that is not impaired by steric hindrance from the rest of the CCG effector. Consistent with this, AlphaFold-Multimer prediction of WRR4A bound to full-length CCG40 indicates a similar binding mode to that of the isolated N-terminal domain, with the C-terminal region projecting away from the receptor-effector interface (Fig. S5B). In the Sr35-Avr35 recognition system, the effector Avr35 exists as a homodimer but dissociates upon binding to Sr35(Forderer et al., 2022). It remains possible that the N-terminal ferredoxin-like domains of CCG effectors engage in loose associations with other domains that dissociate upon recognition by WRR4A.

### Upon activation, the WRR4A resistosome can persist in the absence of effector

The WRR4A-CCG28N structure is notable among plant resistosomes in that it shows little interpretable effector density, resembling a near-apo TIR-NLR resistosome. This observation suggests that, once effector-triggered conformational changes and oligomerization have occurred, the resistosome can remain stable without persistent effector engagement. Such a mechanism would allow NLR activation by effectors with relatively weak or transient binding, as rapid effector dissociation would not necessarily impede resistosome assembly. A common structural feature across all available plant resistosome-effector complexes is that effectors within the resistosomes do not interact with one another or participate in oligomerization interfaces(Wang et al., 2019a; Ma et al., 2020; Martin et al., 2020; Forderer et al., 2022). This indicates that bound effectors in activated NLRs might not play a major role in stabilizing resistosome assembly. The model is supported by the observations that auto-active NLR mutants can form stable oligomeric complexes without any effector binding(Liu et al., 2024; Guo et al., 2025). In the case of WRR4A-CCG28N, after initiating NB domain repositioning, CCG28 can dissociate without impairing resistosome stability. Such released effectors could potentially activate additional WRR4A molecules.

### Resting states of TIR-NLRs with C-JID domains

WRR4A contains a C-terminal C-JID domain characteristic of many TIR-NLRs, which has been shown to contribute to effector recognition(Ma et al., 2020; Martin et al., 2020). However, how C-JID–containing TIR-NLRs maintain autoinhibition prior to activation and how effector binding triggers activation remain poorly understood. Although we were unable to obtain a high-resolution structure of resting state WRR4A, our cryo-EM 2D class averages and AlphaFold predictions converge on a model in which the NB domain occupies the concave surface of the LRR and contacts loops of the C-JID domain. This suggests that the C-JID domain not only mediates effector binding but also contributes to stabilizing the autoinhibited resting state through interactions with the NB domain, potentially representing a shared feature of C-JID–containing TIR-NLRs. Comparison with the resting state structures of the CC-NLRs ZAR1 and NRC2 suggests that the WRR4A LRR domain adopts a more vertical orientation, which may facilitate C-JID–NB interactions that stabilize the inactive state. It will be interesting to see the resting state conformations of TIR-NLRs lacking a C-JID domain, such as flax M(Anderson et al., 1997) and potato Ry_sto(Grech-Baran et al., 2020)_. Despite these architectural differences, the core activation principle appears conserved: the NB domain is constrained in the resting state and displaced upon effector binding to initiate activation.

### Structure-guided engineering of improved resistance

The WRR4A^CCG47^ variant and the WRR4A^W562A^ variant reported here represent two strategies for reprogramming NLR function: creating new recognition capacity and sensitizing receptors. While both strategies have been successfully employed previously using genetic data or random mutagenesis(Segretin et al., 2014; Giannakopoulou et al., 2015), our work demonstrates that structure-guided engineering can accelerate this process, particularly for NLRs with limited genetic information. Together, these results demonstrate that structure-informed design enables rational reprogramming of NLR recognition capacity and strength.

To summarize, our study provides a novel mechanism for broad recognition of multiple effectors by a plant immune receptor, provides insights into the immune receptor resting state, illuminates the activation process of TIR-NLRs with C-JID domains, and demonstrates the potential for extending immune receptor recognition capacity based on structural information.

## ACKNOWLEDGEMENTS

We thank Natasha Lukoyanova and Shu Chen for assistance with cryo-EM imaging at the Birkbeck ISMBEM facility, and Jake Richardson for training in transmission electron microscopy and for maintaining the JIC electron microscopy facility. We are grateful to the Staskawicz laboratory for providing *N. benthamiana eds1* mutant seeds, and to Adam Bentham for assistance during protein purification. We also acknowledge the TSL Bioinformatics Team, particularly Dan MacLean and George Deeks, as well as the JIC Structural Biology Platform, especially David Lawson, for computing support. We thank the TSL Synthetic Biology Team, Media Services, and the Horticultural Team for technical and logistical support. This work was supported by BBSRC grants BB/P021646/1, BB/S018832/1, and BB/W017423/1, and by the Gatsby Charitable Foundation (all awarded to J.D.G.J.). This work was supported by the UKRI Biotechnology and Biological Sciences Research Council Norwich Research Park Biosciences Doctoral Training Partnership [grant number BB/T008717/1]. Cryo-EM data collection was supported by the Wellcome Trust (grants 202679/Z/16/Z and 206166/Z/17/Z). H.Z. was supported by an EMBO Postdoctoral Fellowship (EMBO ALTF88-2021).

## AUTHOR CONTRIBUTIONS

Conceptualization: H.Z., M.S., J.D.G.J; Methodology: H.Z., M.S., J.H.H., F.L.H.M., M.W.W., P.D.; Investigation: H.Z., M.S., C.M.S., X.L. M.W.; Visualization: H.Z, M.S.; Supervision: J.D.G.J, M.S., S.K., Writing–original draft: H.Z., M.S.; Writing–review & editing: H.Z., M.S, J.D.G.J., F.L.H.M., M.W.W., S.K.

