## Supplemental materials for "Structural basis for the broad recognition specificity of an Arabidopsis immune receptor"

Fig. S1

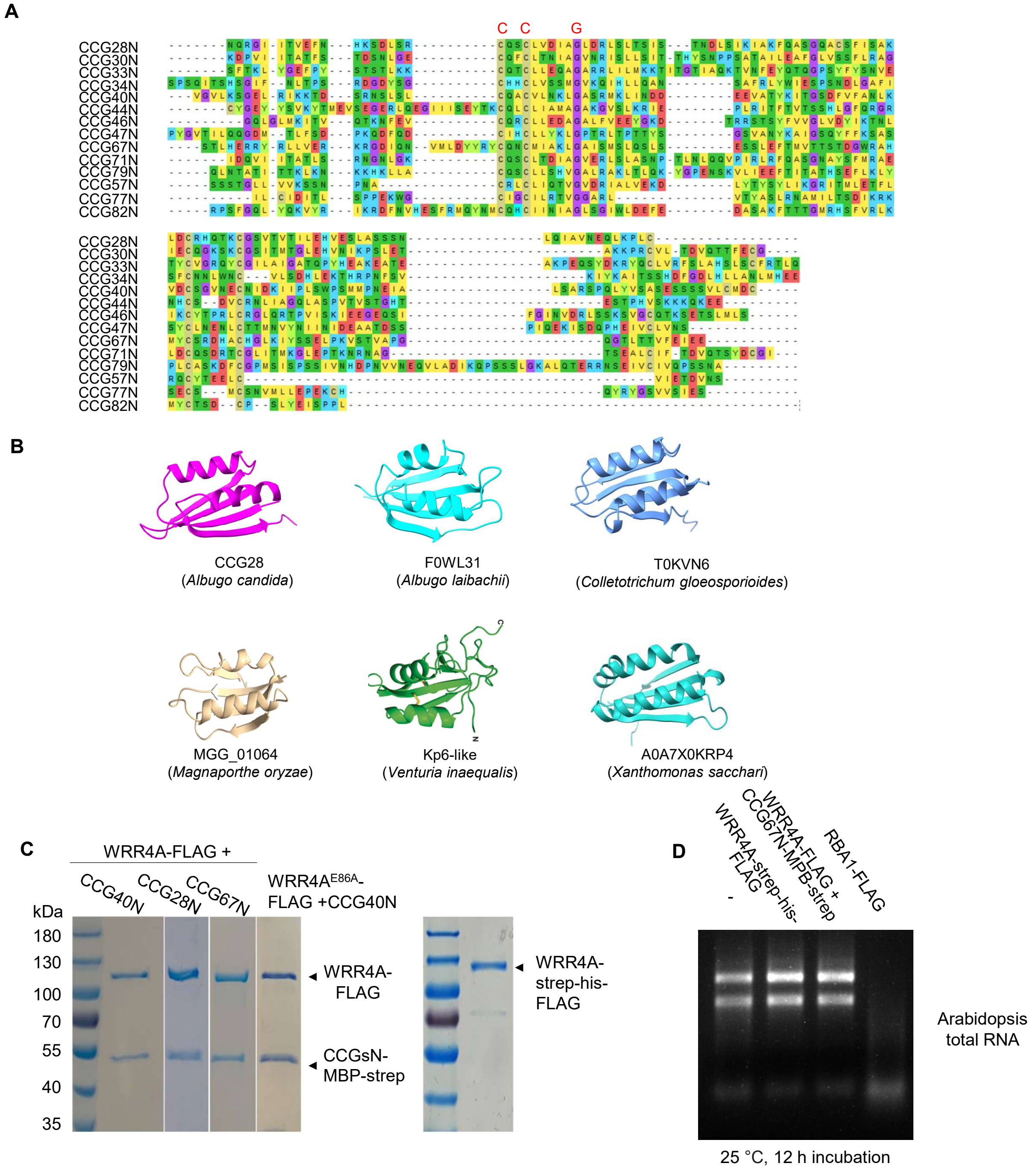


**Fig. S1 CCG and ferredoxin-like effectors**

**(A)** Sequence alignment of N-terminal recognized domains of CCG effectors in Fig. 1A. **(B)** Ferredoxin-like fold effectors from diverse pathogens. CCG N-terminal domains were used as structural queries in Foldseek to identify homologous folds. Candidate proteins were further analyzed with SignalP 6.0 to predict signal peptides. The ferredoxin-like fold domains of candidates were shown. **(C)** Coomassie blue staining of purified protein samples used in this study. Each sample was purified and analyzed separately; 10 μL of each sample was loaded and stained for 15 min before imaging. **(D)** Neither resting-state nor activated WRR4A exhibited visible RNA-hydrolyzing activity. Purified proteins were incubated with 100 ng Arabidopsis total RNA for 12 h at 25 °C before being visualized by agarose gel electrophoresis. The TIR-only protein RBA1 served as a positive control. Image shown is representative of three independent experiments.

Fig. S2

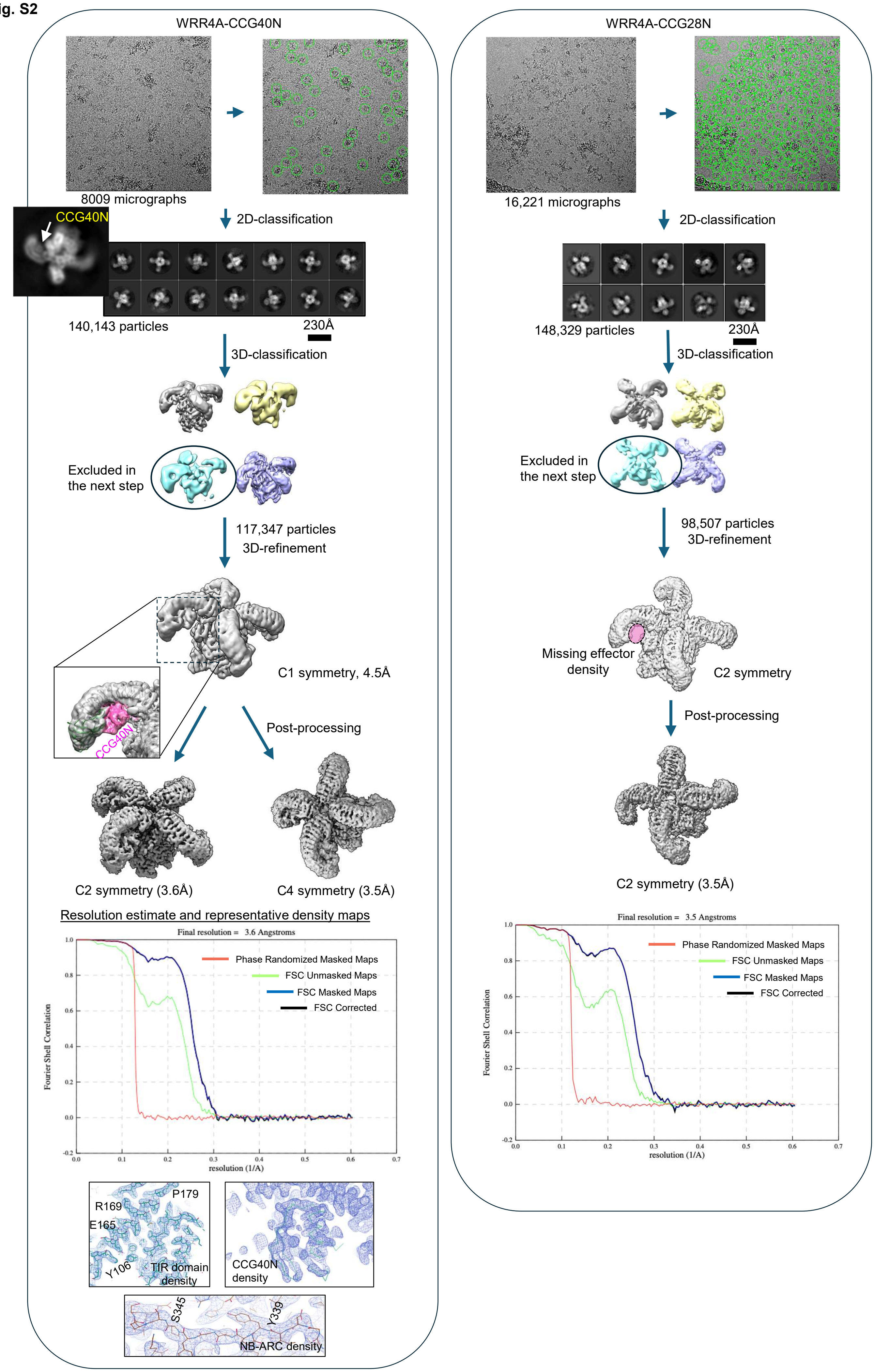

WRR4A-CCG40N
8009 micrographs
CCG40N
2D-classification
140,143 particles
230Å
3D-classification
Excluded in
the next step
117,347 particles
3D-refinement
C1 symmetry, 4.5Å
CCG40N
Post-processing
C2 symmetry (3.6Å)
C4 symmetry (3.5Å)
Resolution estimate and representative density maps
Final resolution = 3.6 Angstroms
Phase Randomized Masked Maps
FSC Unmasked Maps
FSC Masked Maps
FSC Corrected
Fourier Shell Correlation
resolution (1/A)
P179
R169
E165
Y106
TIR domain
density
CCG40N
density
S345
Y339
NB-ARC density
WRR4A-CCG28N
16,221 micrographs
2D-classification
148,329 particles
230Å
3D-classification
Excluded in
the next step
98,507 particles
3D-refinement
Missing effector
density
C2 symmetry
Post-processing
C2 symmetry (3.5Å)
Final resolution = 3.5 Angstroms
Phase Randomized Masked Maps
FSC Unmasked Maps
FSC Masked Maps
FSC Corrected
Fourier Shell Correlation
resolution (1/A)

**Fig. S2 Cryo-EM data processing workflow and 3D reconstruction of WRR4A resistosomes**
Flowchart depicting the cryo-EM data processing pipeline from initial micrograph collection through final 3D reconstruction of WRR4A resistosomes. The number of particles retained at each step is indicated. The Fourier Shell Correlation (FSC) curve (bottom) shows the resolution estimation of the final density map, with the FSC = 0.143 criterion used to determine the overall resolution.

**Fig. S3**

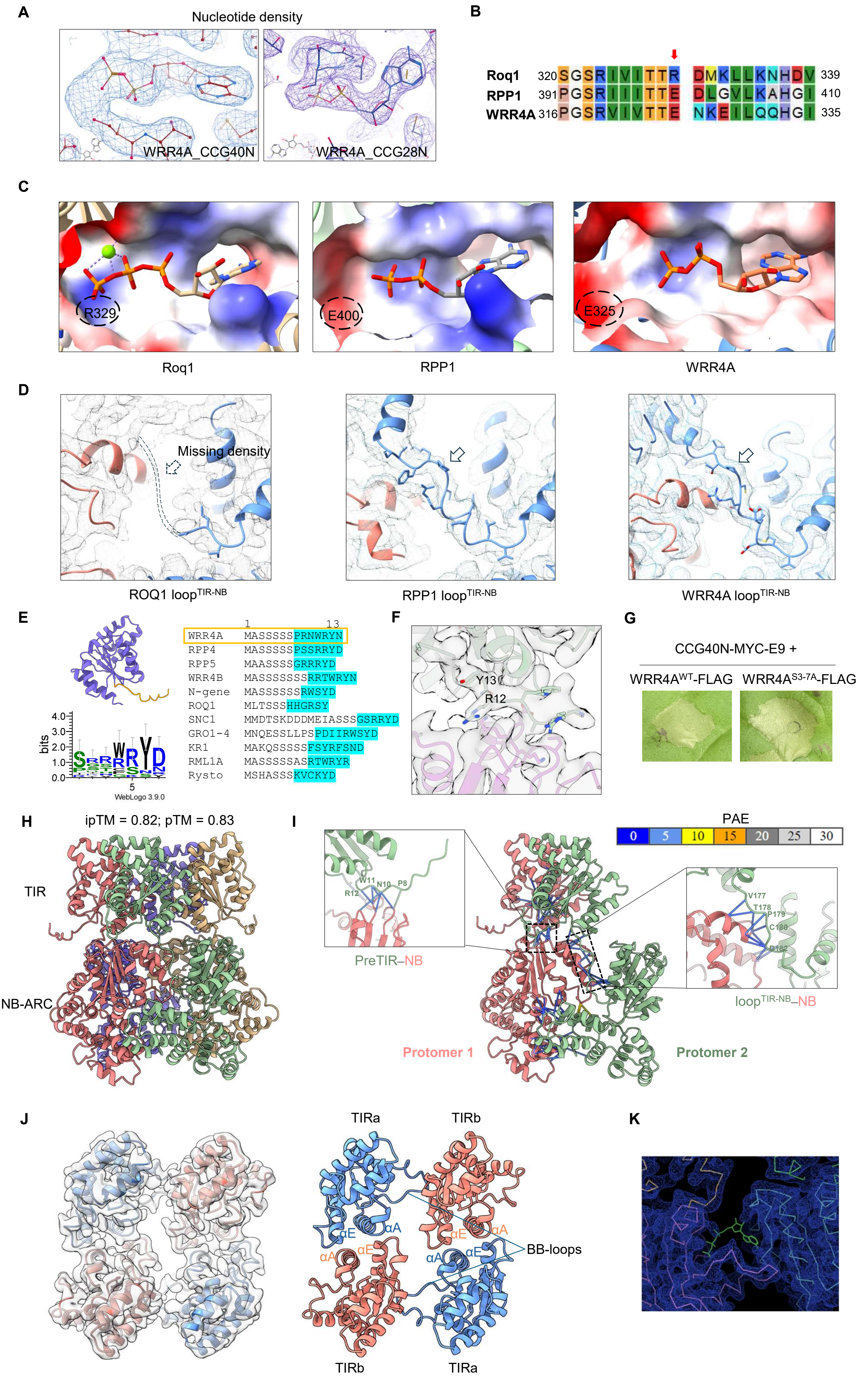
A
Nucleotide density
WRR4A_CCG40N
WRR4A_CCG28N
B
Roq1 320 SGSRIVITTR DMKLLKNHDV 339
RPP1 391 PGSRIIITTE DLGVLKAHGI 410
WRR4A 316 PGSRVIVTTE NKEILQQHGI 335
C
R329
E400
E325
Roq1
RPP1
WRR4A
D
Missing density
ROQ1 loop TIR-NB
RPP1 loop TIR-NB
WRR4A loop TIR-NB
E
1 13
WRR4A MASSSSSPRNWRYN
RPP4 MASSSSSPSSRRYD
RPP5 MAASSSSGRRRYD
WRR4B MASSSSSSSRRTWRYN
N-gene MASSSSSSRWSYD
ROQ1 MLTSSSHHGRSY
SNC1 MMDTSKDDDMEIASSSGSRRYD
GRO1-4 MNQESSLLPSPDIIRWSYD
KR1 MAKQSSSSSFSYRFSND
RML1A MASSSSSASRTWRYR
Rysto MSHASSSKVCKYD
bits
4.0
3.0
2.0
1.0
0.0
5
WebLogo 3.9.0
F
Y13
R12
G
CCG40N-MYC-E9 +
WRR4AWT-FLAG
WRR4AS3-7A-FLAG
H
ipTM = 0.82; pTM = 0.83
TIR
NB-ARC
I
PAE
0 5 10 15 20 25 30
W11
N10
P8
R12
PreTIR–NB
V177
T178
P179
C180
D182
loopTIR-NB–NB
Protomer 1
Protomer 2
J
TIRa
TIRb
αE
αA
αE
αA
αA
αE
αA
αE
BB-loops
TIRb
TIRa
K

**Fig. S3 Structural features of the WRR4A-CCG40 resistosome assembly**

**(A)** Cryo-EM density at the nucleotide-binding site of the WRR4A NB domain indicates ADP binding. **(B)** Sequence alignment of the TT/SR motif region in WRR4A, ROQ1, and RPP1, highlighting the substitution of glutamate with arginine in WRR4A and RPP1. **(C)** Comparison of nucleotide-binding pockets in WRR4A, ROQ1 (PDB ID: 7JLV), and RPP1 (PDB ID: 7CRC). The R-to-E substitution in the conserved TT/SR motif of RPP1 and WRR4A is linked to abolished γ-phosphate coordination. **(D)** Cryo-EM density for the loop$^{TIR\text{-}NB}$ is visible in WRR4A and RPP1 (PDB ID: 7CRC) but absent in ROQ1 (PDB ID: 7JLV). **(E)** Conservation of the pre-TIR motif among TIR-NLRs. The pre-TIR motif position is highlighted in yellow in the TIR domain model. Pre-TIR motif sequences across different TIR-NLRs are shown. Sequence logo of cyan-coloured residues potentially involved in inter-protomer interactions was generated using WebLogo 3 (https://weblogo.threeplusone.com). **(F)** Cryo-EM density for the WRR4A pre-TIR motif showing close contacts with the NB domain of the adjacent protomer. **(G)** Mutation of tandem serine residues to alanine WRR4A$^{S3\text{-}7A}$ in the pre-TIR motif does not affect WRR4A-mediated cell death. Wild-type or mutant WRR4A was transiently co-expressed with CCG40N in *N. tabacum* leaves. Cell death was observed 2 days post-infiltration. Images are representative of three independent replicates. **(H)** AlphaFold3-predicted tetrameric assembly of the WRR4A$^{TIR\text{–}NB\text{-}ARC}$ module. **(I)** Close-up view of the inter-protomer interactions from (H). Interactions mediated by the loop$^{TIR\text{-}NB}$ and the pre-TIR motif are highlighted. Solid lines indicate residue contacts within 4 Å, coloured by predicted aligned error (PAE) from AlphaFold using ChimeraX. **(J)** The TIR domains in the WRR4A resistosome adopt a dimer-of-heterodimer conformation. **(K)** No detectable cryo-EM density at the substrate-binding site of the WRR4A TIR heterodimer.

Fig. S4

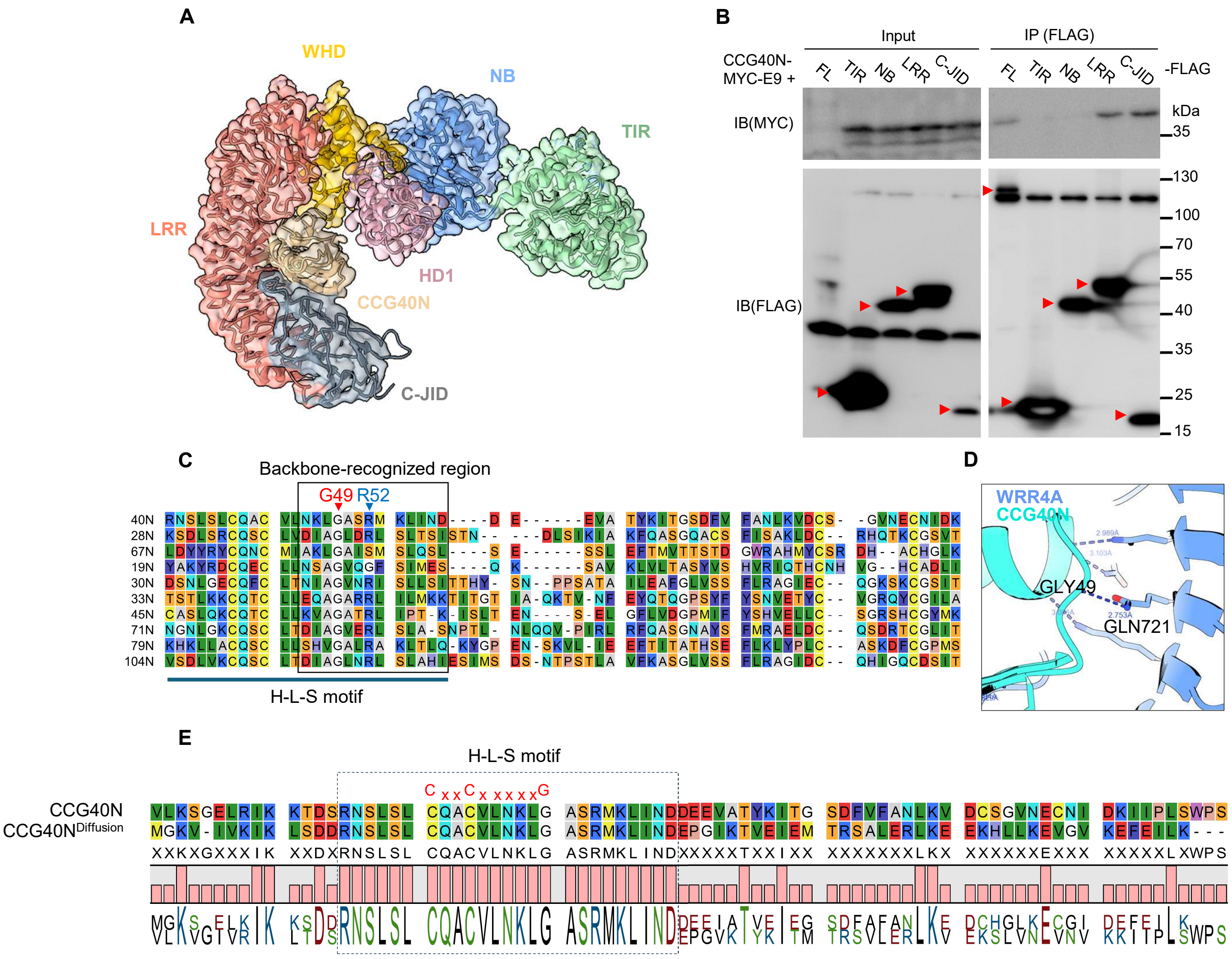


**Fig. S4 Recognition of CCG40N by WRR4A**

**(A)** Cryo-EM density of a protomer from the WRR4A-CCG40N complex, showing domain architecture of activated WRR4A in complex with the effector. **(B)** WRR4A interacts with CCG40N primarily through its LRR and C-JID domains. Individual FLAG-tagged WRR4A domains were co-expressed with CCG40N-MYC-E9 in *N. benthamiana eds1* mutant leaves. Total protein were immunoprecipitated with anti-FLAG beads and detected with anti-FLAG and anti-MYC antibodies. FL indicates full length WRR4A. **(C)** The H-L-S motif and its backbone-recognition region among CCG effectors. The conserved "G" in the CCG motif and a relatively conserved R involved in recognition are highlighted. **(D)** Interaction between the conserved "G" in the CCG motif of CCG40N and the side chain of WRR4A Gln721. **(E)** Sequence alignment of CCG40N and CCG40N$^{\text{Diffusion}}$. The retained H-L-S motif during diffusion is indicated.

Fig. S5

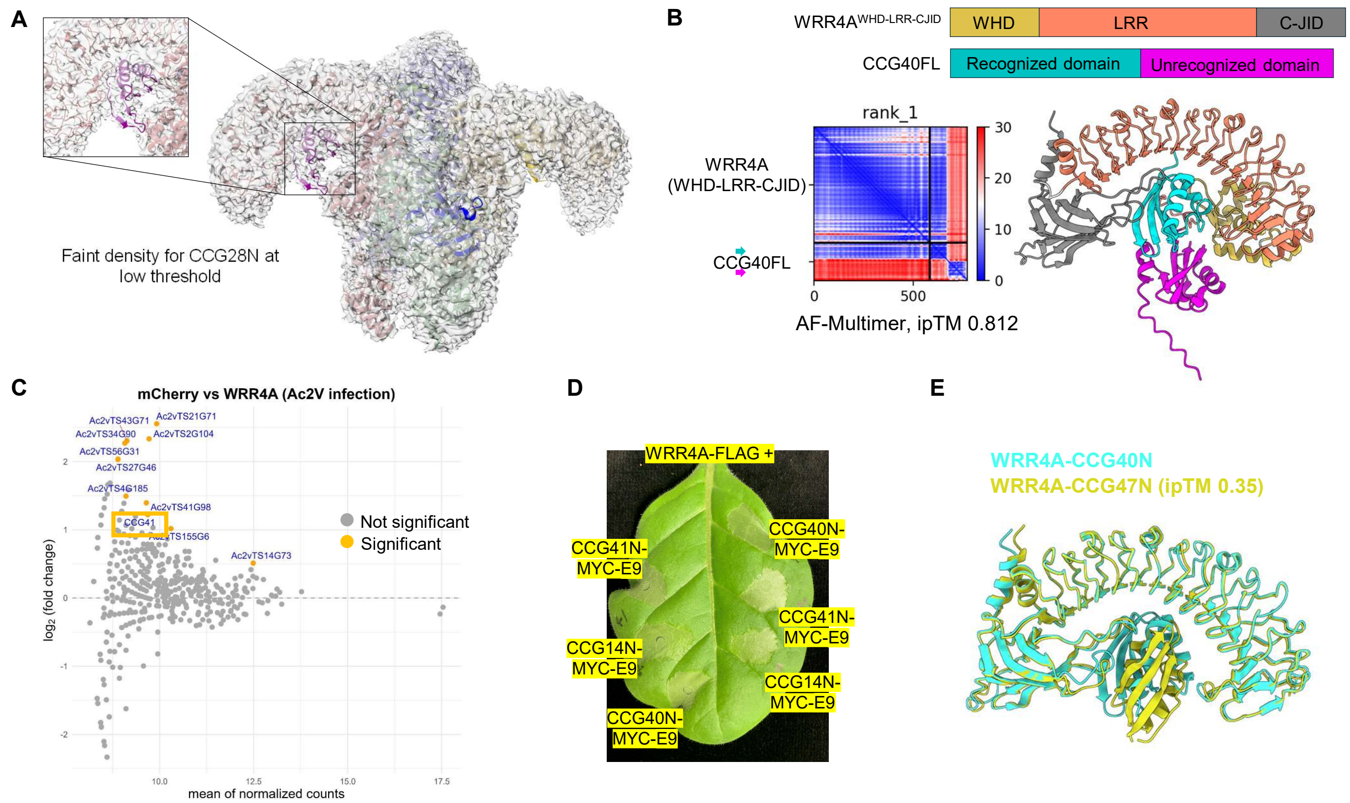


F

| WRR4A | CCG40N | Corresponding residues in CCG47N | WRR4A$^{CCG47}$ |
|---|---|---|---|
| TYR979 | LYS30 | SER16 | |
| GLN900 | ARG34 | GLN20 | |
| ASP981 | ARG34 | GLN20 | Asn |
| GLU903 | ASN35 | ASP21 | Lys |
| THR742 | SER51 | THR36 | |
| ASP788 | ARG52 | ARG37 | |
| GLU831 | LYS54 | THR39 | |
| TYR653 | ASN76 | SER61 | |
| ASN700 | ASN46 | TYR31 | |
| ASN702 | ASN46 | TYR31 | |
| LYS703 | ASN46 | TYR31 | |
| GLN721 | GLY49 | GLY34 | |
| LYS744 | ASN46 | TYR31 | Ala |
| ASN908 | ASP58 | TYR43 | |

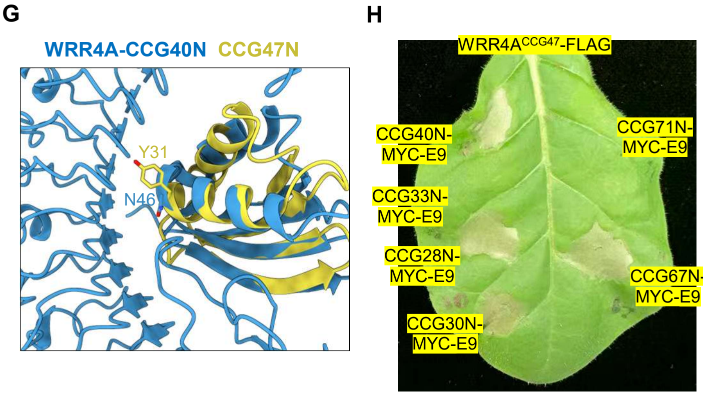


**Fig. S5 Accurate AlphaFold-Multimer predictions support WRR4A recognition engineering**
**(A)** Cryo-EM density map of the WRR4A-CCG28 complex. Faint density is visible at the effector-binding site at low contour threshold, facilitating rigid-body fitting of CCG28N. **(B)** AlphaFold-Multimer prediction of WRR4A in complex with full-length CCG40. The PAE plot indicates that the unrecognized C-terminal domain of CCG40 cannot be confidently positioned relative to either CCG40 recognized domain or WRR4A domains, suggesting potential flexibility. **(C)** TurboID-based identification of CCG14 (Ac2vTS47G216) and CCG41 (Ac2vTS3G55) as WRR4A-recognized effectors. WRR4A-TurboID transgenic Arabidopsis *eds1* mutants were infected with *A. candida* for 6 days, followed by biotin labelling and streptavidin IP-MS. CCG41 was significantly enriched compared to the mCherry-TurboID control (log2(FC) > 0 and P-value < 0.05). CCG14 was detected exclusively in WRR4A-TurboID samples but not controls. **(D)** WRR4A recognizes the N-terminal regions of CCG14 and CCG41. WRR4A was transiently co-expressed with CCG14N or CCG41N in *N. tabacum* leaves. Cell death was assessed 2 days post-infiltration. **(E)** The AlphaFold-Multimer model of WRR4A-CCG47N shows low ipTM score and does not match the binding mode observed for CCG40N. **(F)** Polar interactions between WRR4A and CCG40N, and the corresponding amino acid substitutions in CCG47N. **(G)** Tyrosine substitution in CCG47 at the position corresponding to glutamine in CCG40. **(H)** Recognition specificity of WRR4A$^{CCG47}$. WRR4A$^{CCG47}$ was transiently co-expressed with different CCGs in *N. tabacum leaves*. Cell death was assessed 2 days post-infiltration.

Fig. S6

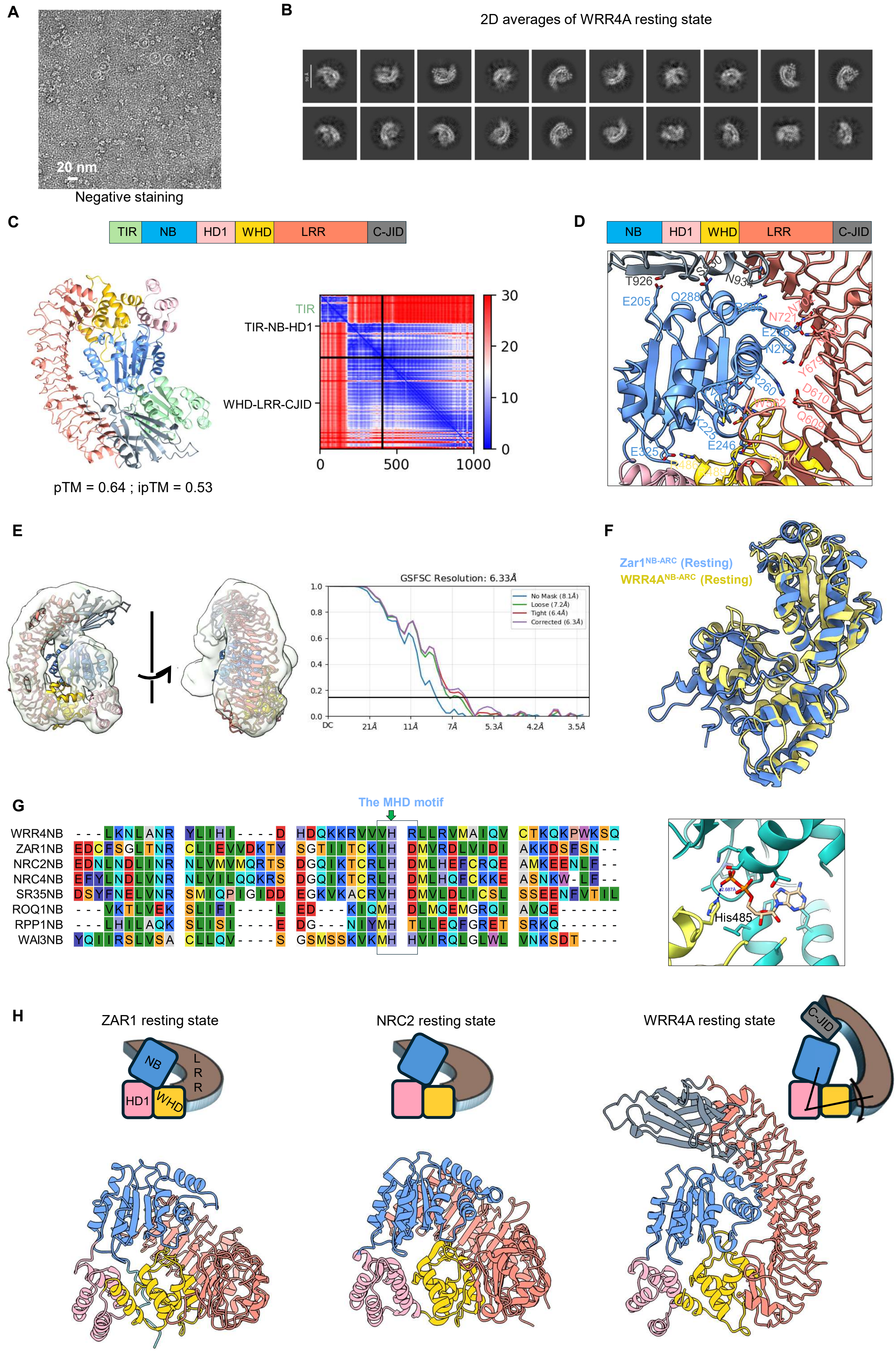
A
20 nm
Negative staining
B
2D averages of WRR4A resting state
C
TIR
NB
HD1
WHD
LRR
C-JID
TIR
TIR-NB-HD1
WHD-LRR-CJID
30
20
10
0
0
500
1000
pTM = 0.64 ; ipTM = 0.53
D
NB
HD1
WHD
LRR
C-JID
T926
E205
Q288
N934
N721
E246
E325
D610
Q609
E
GSFSC Resolution: 6.33Å
No Mask (8.1Å)
Loose (7.2Å)
Tight (6.4Å)
Corrected (6.3Å)
DC
21Å
11Å
7Å
5.3Å
4.2Å
3.5Å
F
Zar1NB-ARC (Resting)
WRR4ANB-ARC (Resting)
G
The MHD motif
WRR4NB
ZAR1NB
NRC2NB
NRC4NB
SR35NB
ROQ1NB
RPP1NB
WAI3NB
His485
H
ZAR1 resting state
NRC2 resting state
WRR4A resting state
NB
LRR
HD1
WHD
C-JID

**Fig. S6 Structural analysis of resting-state WRR4A**

**(A)** Negative-stain electron microscopy of resting state WRR4A. **(B)** 2D class averages of cryo-EM data of WRR4A resting state showing monomeric architecture. **(C)** AlphaFold-Multimer prediction of full-length WRR4A resting state modelled as two chains (TIR-NB-HD1 and WHD-LRR-C-JID). The PAE plot indicates TIR domain positioning relative to other domains cannot be confidently predicted. **(D)** Interactions between the WRR4A NB domain and the WHD, LRR, and C-JID domains in the resting state. **(E)** Low-resolution 3D reconstruction of resting-state WRR4A showing absence of interpretable TIR domain density. **(F)** The NB-HD1-WHD domain arrangement in predicted resting-state WRR4A resembles ZAR1 (PDB ID: 6J5T). **(G)** The MHD motif is accurately modelled with the conserved histidine positioned to coordinate the β-phosphate of ADP. **(H)** Comparison of resting-state NLR structures reveals distinct domain arrangements. In WRR4A, the NB domain lies within the concave curvature of the LRR, surrounded by WHD, LRR, and C-JID domains. The WRR4A LRR adopts a vertical orientation, lying approximately in the plane of the NB-HD1-WHD domains, contrasting with ZAR1 (PDB ID: 6J5T) and NRC2 (PDB ID: 8RFH).

**Supplementary Table 1 List of potential interacting residues between WRR4A and CCG40N or CCG28N**

| Structure | WRR4A | Effector |
|---|---|---|
| **WRR4A-CCG40N** | R376, Y442, E443, R481, A560, R633, Y653, H675, M677, Y679, F698, N700, N702, K703, D719, Q721, T742, K744, H764, C790, F902, P906, N908, H928, S930, H931, M932, F933, N934, A935, Y979 | R34, Q41, N46, L47, L48, G49, A50, S51, R52, M53, K54, L55, I56, N57, D58, S70, F72, V73, A75, K78, G83, V84, D90 |
| **WRR4A-CCG28N** | D365, M368, D369, Y442, Q478, K479, R481, A560, G561, W562, H563, S632, R633, Y653, H675, Y679, F698, N700, N702, Y703, D719, Q721, H787, L904, P906, H928, H931, F933, N934, A935, Y979 | S45, Q48, S49, V52, D53, I54, D58, L60, L62, I65, Q68, D69, L70, Q82, C84, S85, A83, I87, S88, K90, D92, H95, Q96, T97, K98, T105, I106, |

**Supplementary Table 2 Details of cryo-EM structural analyses in this study**

| **Data collection parameters** | WRR4A-CCG40N<br>PDB ID 9QU9, EMD-53375 | WRR4A-CCG28N<br>PDB ID 9R8W, EMD-53844 | Resting state WRR4A |
|---|---|---|---|
| Microscope | Titan Krios D3771 | Titan Krios D3771 | Titan Krios D3771 |
| Acceleration voltage (kV) | 300 | 300 | 300 |
| Pixel size | Gaton K3 | Gaton K3 | Gaton K3 |
| Magnification | 105,000 | 105,000 | 105,000 |
| Detector | 0.828 | 0.828 | 0.828 |
| Defocus range (μm) | -2.7 to -1.6 | -2.7 to -1.6 | -2.7 to -1.6 |
| Frames per exposure | 50 | 50 | 50 |
| Electron dose | 50e/Å$^2$ | 50e/Å$^2$ | 50e/Å$^2$ |
| Exposure per hole | 2 | 4 | 4 |
| Data collection software | EPU3.4 | EPU3.4 | EPU3.4 |
| Number of Micrographs | 8009 | 16,211 | 7,497 |
| Final refined particles | 117,347 | 98,507 | - |
| Symmetry imposed | C2 | C2 | - |
| Unmasked resolution at 0.143 cut-off FSC (Å) | 3.9 | | - |
| Masked resolution at 0.143 cut-off FSC (Å) | 3.6 | 3.5 | - |
| Map shaprening B factor (Å$^2$) | 142 | 50 | - |
| **Model building and Refinement** | | | |
| <u>Model composition</u> | | | - |
| Protein | 4 chains of WRR<br>4 chains of effector | 4 chains of WRR<br>4 chains of effector | - |

| | | | |
|---|---|---|---|
| Ligands | 4 molecules of ADP | 4 molecules of ADP | - |
| Validation | | | - |
| RMSD Bond (Å) | 0.003 | 0.003 | - |
| RMSD angles (°) | 0.761 | 0.649 | - |
| MolProbity score | 2.39 | 2.54 | - |
| Clash score | 13.57 | 13.34 | - |
| Rotamer outliers (%) | 2.42 | 3.24 | - |
| C-beta outliers (%) | 0.0 | 0.0 | - |
| CC (mask) | 0.86 | 0.81 | - |
| CC (volume) | 0.83 | 0.79 | - |
| EMRinger | 2.38 | 1.44 | - |
| Ramachandrann plot | | | |
| Favoured (%) | 92.67 | 91.17 | - |
| Allowed (%) | 6.89 | 8.57 | - |
| Outliers (%) | 0.43 | 0.26 | - |

### Supplementary Table 3 Primers used in the study

| Primer | Sequence | Purpose |
|---|---|---|
| WRR4A-F | GCGGTCTCAAATGGCTTCTTCTTCTTCCTC | Full length WRR4A cloning |
| WRR4A-R | GCGGTCTCACGAACCGAATACTAGAGAACCCCAAA | Full length WRR4A cloning |
| WRR4A-534-F | GC GGTCTC A AATG CCGTGTCGGGATTTTGATGAC | Domain mapping |
| WRR4A-534-R | GC GGTCTC A CGAA CC TGTGACATTTATTTTAGTCGAAACAT | Domain mapping |
| WRR4-1561-F | GC GGTCTC A AATG ACGGGTAATCGATCTATTAAAGG | Domain mapping |
| WRR4-2602-F | GC GGTCTC A AATG GAGTTCGGCCACCGAGCC | Domain mapping |
| WRR4-1560-R | GC GGTCTC A CGAA CC TGCTTCTTCAAGAACATAAGCTATCT | Domain mapping |
| WRR4-2601-R | GC GGTCTC A CGAA CC TGGAGGTACTTCTAATCCTGGT | Domain mapping |
| WRR4A_N176A_T178A_R181A_D182A-F1 | GCGGTCTCAATGTTTCGACTAAAATAGCTGTCGCACCGTGTGCGGCTTTTGATGACATGGT | Point mutation |
| WRR_N176A_T178A_R181A_D182A-R1 | GCGGTCTCAACATCTCTTGCTATCTTCTCTATCA | Point mutation |
| WRR_D182I_F1 | GCGGTCTCACACCGTGTCGGATTTTTGATGACA | Point mutation |
| WRR_D182I_R1 | GCGGTCTCAGGTGTGACATTTATTTTAGTCGAAA | Point mutation |
| WRR_preTIR_F: | GCGGTCTCAAATGGCTTCTTCTTCTTCCTCATCTGCCGCCTCGGCATCCAATGTCTTCACGAGCTTCCAT | Point mutation |
| HZ_WRR4-CDS-K703E-F1 | GC GGTCTC A CAACATGAATGAATGCTCAAGAT | Point mutation |
| HZ_WRR4-CDS-K703E-R1 | GC GGTCTC A GTTGAGGAAGGTAAGAGATGTCA | Point mutation |
| HZ_WRR4A_148A_R1 | GCGGTCTCATCGTAGACTTTCAGGAACAAAAG | Point mutation |
| HZ_WRR4A_W148A_F1 | GCGGTCTCAACGATGCGGGGGCGCATACCGG | Point mutation |
| HZ_WRR4A-S930P-F1 | GCGGTCTCATCACCTTCCACATATGTTCAACG | Point mutation |
| HZ_WRR4A-S930P-R1 | GCGGTCTCAGTGAAACGTCTTTTCATCAGAG | Point mutation |
| WRR4nd1-SWAP-F | GCGGTCTCAGATAGCTTATGTTCTTGAAGAAGCA | Nd1LRRswap |
| WRR4nd1-SWAP-R | GCGGTCTCATATCTTTTCGGCATCTACTAGAATC | Nd1LRRswap |
| WRR4A-S5A5-F | GCGGTCTCAAATGGCTGCTGCTGCTGCTGCTCCTCGCAACTGGAGATACAATGTCTTCACG | PreTIR-S5A5 |
| HZ_WRR4A-D788A-F1 | GCGGTCTCATAATCTACACGCCCTTTGTCTA | Point mutation |
| HZ_WRR4A-D788A-R1 | GCGGTCTCAATTATGAAGACCTTTGATGCAATCT | Point mutation |
| CCG33-31-F | GC GGTCTC A AATG GGTGAATTTCCGTACAGCAC | CCG33N cloning |
| CCG33-129-R | GC GGTCTC A CGAA CC CTGATACCGCTTGTCATAGC | CCG33N cloning |
| CCG40-20-F | GC GGTCTC A AATG GTTCTAAAATCAGGCGAGTTGAG | CCG40N cloning |
| CCG40-130-R | GC GGTCTC A CGAA CC ACAGTCCATGCACAAGACAC | CCG40N cloning |
| CCG67-23-F | GC GGTCTC A AATG AGTACACTTCACGAAAGAAGATACC | CCG67N cloning |
| CCG67-130-R | GC GGTCTC A CGAA CC TTCTTCGATTTCAAAAACAGTCG | CCG67N cloning |
| CCG79-17-F | GC GGTCTC A AATG TGTCATATCCACCAAGCGTA | CCG79N cloning |
| CCG79-130-R | GC GGTCTC A CGAA CC ACTCGAGGGTTGCTTGATGT | CCG79N cloning |
| CCG104-22-F | GC GGTCTC A AATG TCCTGCCTAACAGACATTGC | CCG104N cloning |
| CCG104-130-R | GC GGTCTC A CGAA CC AGAGCCACTCGCCATTTTAA | CCG104N cloning |
| HZ_CCG34N_F | GCGGTCTCAAATGTCGCCCAGTCAGATAACCTC | CCG34N cloning |
| HZ_CCG34N_R | GCGGTCTCACGAACCCTCTTCATGCATCAAGTTCG | CCG34N cloning |
| HZ_CCG46N_F | GCGGTCTCAAATGGGTCAGCTGGGCTTGATG | CCG46N cloning |
| HZ_CCG46N_R | GCGGTCTCACGAACCGCTAAGCATTAAAGAAGTCTC | CCG46N cloning |
| HZ_CCG40GtoW-F1 | GCGGTCTCACAACAAACTGTGGGCTTCCCGA | CCG40 G mutation |
| HZ_CCG40GtoL-F1 | GCGGTCTCACAACAAACTGTTGGCTTCCCGA | CCG40 G mutation |
| HZ_CCG40GtoQ-F1 | GCGGTCTCACAACAAACTGCAGGCTTCCCGA | CCG40 G mutation |
| HZ_CCG40GtoK-F1 | GCGGTCTCACAACAAACTGAAGGCTTCCCGA | CCG40 G mutation |
| HZ_CCG40GtoS-F1 | GCGGTCTCACAACAAACTGAGCGCTTCCCGA | CCG40 G mutation |
| HZ_CCG40GtoD-F1 | GCGGTCTCACAACAAACTGGACGCTTCCCGA | CCG40 G mutation |
